# Donut PCR: a rapid, portable, multiplexed, and quantitative DNA detection platform with single-nucleotide specificity

**DOI:** 10.1101/2020.04.24.058453

**Authors:** Dmitriy Khodakov, Jiaming Li, Jinny X. Zhang, David Yu Zhang

## Abstract

Current platforms for molecular analysis of DNA markers are either limited in multiplexing (qPCR, isothermal amplification), turnaround time (microarrays, NGS), quantitation accuracy (isothermal amplification, microarray, nanopore sequencing), or specificity against single-nucleotide differences (microarrays, nanopore sequencing). Here, we present the Donut PCR platform that features high multiplexing, rapid turnaround times, single nucleotide discrimination, and precise quantitation of DNA targets in a portable, affordable, and battery-powered instrument using closed consumables that minimize contamination. We built a bread-board instrument prototype and three assays/chips to demonstrate the capabilities of Donut PCR: (1) a 9-plex mammal identification panel, (2) a 15-plex bacterial identification panel, and (3) a 30-plex human SNP genotyping assay. The limit of detection of the platform is under 10 genomic copies in under 30 minutes, and the quantitative dynamic range is at least 4 logs. We envision that this platform would be useful for a variety of applications where rapid and highly multiplexed nucleic acid detection is needed at the point of care.

---

DNA and RNA sequence information uniquely identify biological organisms, from human to microbe. Consequently, detection of specific DNA sequences has become a critical part of precision medicine, from pathogen identification to human genetic disease risk assessment to disease prognosis. It is evident that as our understanding of disease genomics improves, translation of this scientific knowledge into actionable clinical practice will be facilitated by DNA diagnostic platforms that are simultaneously fast, affordable, sensitive, massively multiplexed, quantitative, and easy to operate.

Since the early 2000’s, however, DNA detection technologies have bifurcated into either massively multiplexed but slow platforms (NGS [1, 2] and microarrays [3, 4]) or rapid but low multiplexing platforms (qPCR [7, 8] and isothermal amplification [9, 10]). Two notable exceptions to the slow but powerful or fast but limited tradeoff are the Biofire FilmArray multiplex PCR platform [11] and the Oxford Nanopore high-throughput sequencing platform [12] (Table 1). However, both platforms are unable to quantitate accurately and unable to reliably recognize single nucleotide difference, that serve as critical DNA biomarkers for genetic and metabolic risk assessment [13, 14], pharmacogenetic drug dosing [15], cancer therapy selection [16], and infectious disease antimicrobial resistance [17, 18].

**Table 1.**
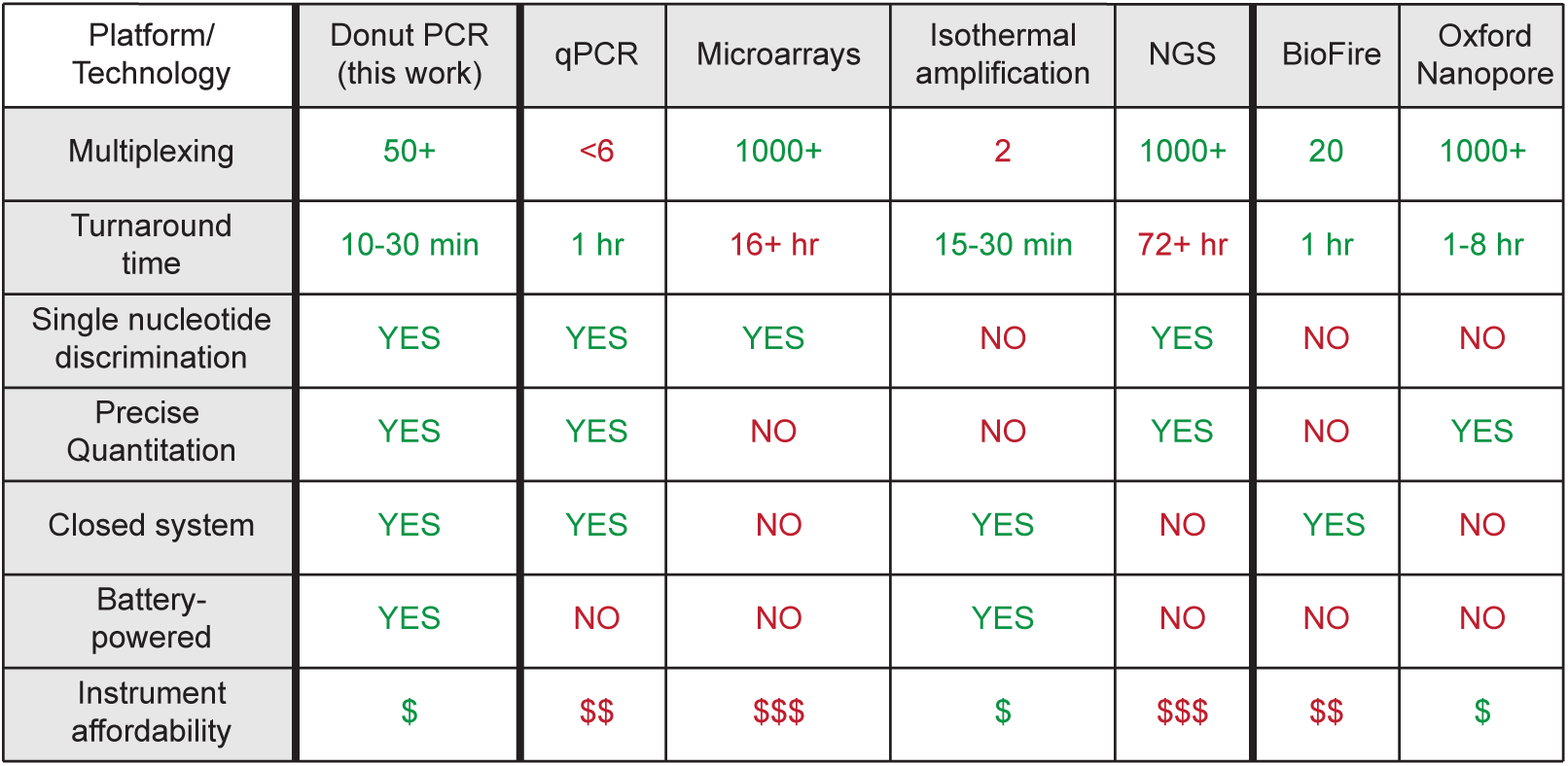
Specification comparison of the existing nucleic acid analysis platforms.

| Platform/<br>Technology | Donut PCR<br>(this work) | qPCR | Microarrays | Isothermal<br>amplification | NGS | BioFire | Oxford<br>Nanopore |
| --- | --- | --- | --- | --- | --- | --- | --- |
| Multiplexing | 50+ | <6 | 1000+ | 2 | 1000+ | 20 | 1000+ |
| Turnaround<br>time | 10-30 min | 1 hr | 16+ hr | 15-30 min | 72+ hr | 1 hr | 1-8 hr |
| Single nucleotide<br>discrimination | YES | YES | YES | NO | YES | NO | NO |
| Precise<br>Quantitation | YES | YES | NO | NO | YES | NO | YES |
| Closed system | YES | YES | NO | YES | NO | YES | NO |
| Battery-powered | YES | NO | NO | YES | NO | NO | NO |
| Instrument<br>affordability | \$ | \$\$ | \$\$\$ | \$ | \$\$\$ | \$\$ | \$ |

Here, we present the Donut PCR platform for DNA detection that combines scalable and massive multiplexing, rapid turnaround times, single nucleotide discrimination, and precise quantitation in a portable, affordable, and battery-powered instrument using closed consumables that minimize contamination risks (Table 1). The Donut PCR system is enabled by two inventions: (1) reliable convection PCR using an annular reaction chamber, and (2) a pre-quenched microarray that allows multiplexed readout via spatial separation. Convection PCR achieves thermal cycling of a PCR reaction mixture using passive movement of fluid due to temperature-induced density differences, enabling an affordable, portable, low-power, and rapid turnaround instrument. The pre-quenched microarray uses spatial separation of fluorescent probes that become unquenched upon hybridization by DNA amplicons, enabling scalable multiplex readouts without open-tube wash steps.

Importantly, the pre-quenched microarray is integrated in the consumable allowing probe hybridization to occur concurrently with convection PCR amplification. Consequently, unlike standard DNA microarrays that visualize spot end-point fluorescence after 16 hours of hybridization, the Donut PCR platform performs real-time detection and quantitation of DNA in under 30 minutes. The quantitative dynamic range is over 4 logs, with a limit of detection of under 10 genomic DNA copies. Use of toehold probes [19] or X-probes [20] for the microarray further provides single nucleotide discrimination that enables robust single nucleotide polymorphism (SNP) genotyping.

## Results

### Donut PCR mechanism and chip design

Rayleigh-Benard thermal convection is the physical principle that as an aqueous solution is heated, it becomes less dense and rises due to gravity, whereas, in contrast, a colder solution is denser and falls. In the Donut PCR platform, we designed a chip that includes an annular (donut-shaped) reaction chamber in which the DNA sample and PCR reagents are loaded. The chip is then sealed and vertically mounted on one side to a 95 °C heater and on the other side to a 60 °C heater (Fig. 1ab). Fluid in the reaction chamber is heated at the 95°C zone and rises to the top of the chamber where it is carried by momentum to the 60°C zone. The fluid is then cooled at the 60°C zone, and falls to the bottom of the chamber, where it is carried by momentum to the 95°C zone, thus completing the thermal cycle.

**FIG. 1:**
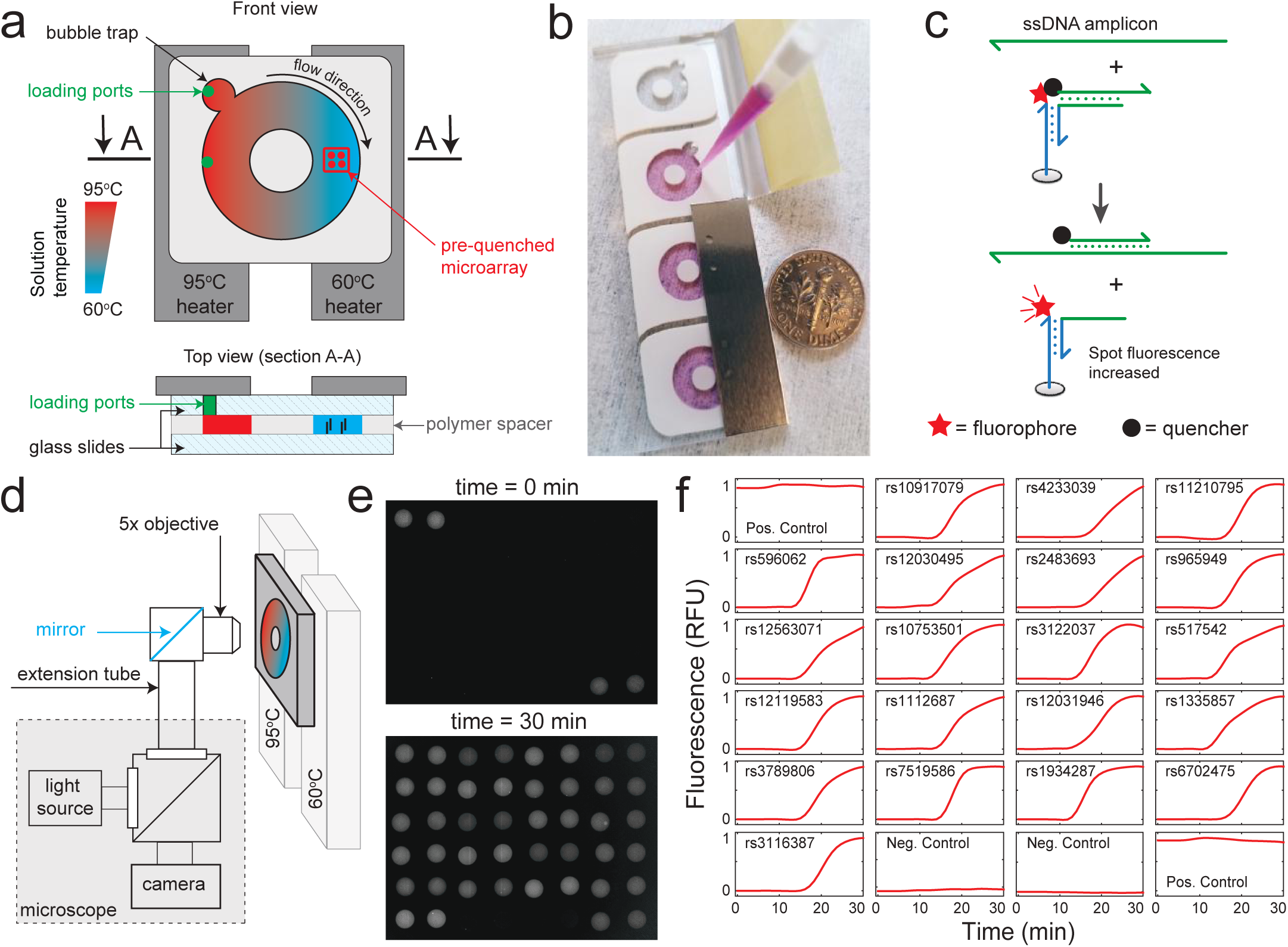
Core innovations in the Donut PCR platform. **(a)** Thermal cycling based on passive fluidic flow, using Rayleigh-Benard convection. PCR reaction mixture comprising DNA template, primers, DNA polymerase, and dNTPs are loaded via the bottom green loading port into the donut-shaped reaction chamber on the Donut PCR chip. Subsequently, the left side of the chip is heated to 95 °C and the right side to 60 °C. Because water is less dense at higher temperatures, the reaction solution will rise on the left side as it is heated and fall on the right side as it is cooled, achieving autonomous thermal cycles with only two constant-temperature heaters. For each lap around the donut-shaped “racetrack”, amplicons have a chance to hybridize to the surface-functionalized probe array printed in the 60 °C zone. **(b)** Picture of a solution of food coloring being loaded into the Donut PCR chip with a US dime for size comparison. **(c)** Schematic of pre-quenched microarray probes printed in the 60 °C zone. At each spot, a three-stranded DNA complex is printed in which the fluorophore-functionalized oligo is colocalized to a quencher-labeled oligo via an unmodified bridge oligo. Green regions show gene-specific DNA sequences and blue regions show universal DNA sequences. The double-stranded DNA (dsDNA) amplicon is denatured in the 95 °C zone to become single-stranded DNA (ssDNA). The ssDNA amplicon then binds to and displaces the quencher-labeled oligo from the surface probe, resulting in a local fluorescence increase. **(d)** Overview of initial chip imaging setup using Zeiss Axiovert 1M epifluorescence microscope. **(e)** Fluorescence image of chip microarray zone before (top) and after (bottom) PCR amplification. The field of view is ≈2 mm wide and each spot has diameter ≈150 *μ*m. Here, probes are printed as adjacent duplicate pairs. The top left and bottom right pair are positive controls, and the bottom middle pairs are negative controls. The remaining 20 pairs of probes target different single-copy regions of the human genome. 10 ng of NA18537 human genomic DNA (roughly 3000 haploid genome copies) was used as input for this experiment, and 20 pairs of corresponding PCR primers were introduced in the reaction to simultaneously amplify all loci of interest within the Donut PCR chip. **(f)** Time-based spot fluorescence for the experiment with endpoint is shown in panel (e). Plotted is the averaged relative fluorescence from each pair of spots. The rs numbers in each subfigure indicate the human single nucleotide polymorphism (SNP) marker included within each amplicon region. Our probes in this experiment only detect the presence of the amplicon and not specific SNP alleles (see also Fig. 5).

On the inner surface of the reaction chamber of the chip, we print a pre-quenched DNA microarray (Fig. 1c) to allow highly multiplexed probe-based readout. Microarrays allow detection of up to hundreds of thousands of different nucleic acid targets using a single fluorescence channel [5, 6] by spatially separating different probes. However, traditional microarrays are unsuitable for in vitro diagnostic (IVD) use because they require labor-intensive and open-tube wash steps to suppress fluorescence background. The team has developed a new pre-quenched microarray chemistry, in which unlabeled amplicons induce an increase of the corresponding spot’s fluorescence via displacement of a quencher-functionalized oligonucleotide. Because the PCR amplicons are unlabeled, no wash steps are needed to reduce fluorescent background from excess amplicons, and a highly multiplexed readout for many different DNA targets can be achieved in a closed tube reaction without specialized opto-fluidic equipment.

Once the loaded chip is mounted against the heaters, the PCR reaction begins, and an external camera is used to periodically take pictures of the microarray area (Fig. 1d). At early time points, probe spots on the microarray are dark except for positive control spots (Fig. 1e); at the end of the reaction, the probes that are hybridized to DNA amplicons become bright. Importantly, in the Donut PCR system, PCR amplification occurs concurrently with probe hybridization, so the progress of the amplification reaction can be tracked in real time, unlike standard microarrays (Fig. 1f). This real-time readout allows robust and accurate DNA target quantitation based on the time at which the fluorescence significantly increases. In contrast, endpoint fluorescence quantitation for standard microarrays is known to be less stable due to sample contents, probe synthesis impurities, illumination non-uniformity, optical aberrance, physical smudges, and other factors.

### Specificity, speed, sensitivity, and quantitation dynamic range

Convection PCR was initially proposed and experimentally demonstrated in 2002 [21], using capillary tubes that are heated from the bottom. However, previous implementations of convection PCR have not entered mainstream use because they exhibited poor temperature uniformity that resulted in significant primer dimer formation and nonspecific genomic amplification. Thus, one of the first priorities in building a convection-based massively multiplex qPCR platform is to ensure PCR amplification specificity.

Based on our understanding, the fluid circulation in a reaction tube or chamber adopts laminar flow, with the flow velocity dependent on the temperature differential of the different circuit paths. In many reaction chamber designs, there will be regions containing fluid paths with minimal temperature differential that have minimal fluid movement (Supplementary Section S5). These regions may result in disproportionate formation of primer dimers and nonspecific amplification. By engineering a donut-shaped reaction chamber in the PCR chip, we remove most of the dead volume, and are able to achieve similar PCR specificity on human genomic DNA as the commercial Bio-Rad CFX96 instrument (Fig. 2a). In contrast, a “pizza” shaped reaction chamber and a capillary tube both result in significant primer dimer and nonspecific amplicon formation.

**FIG. 2:**
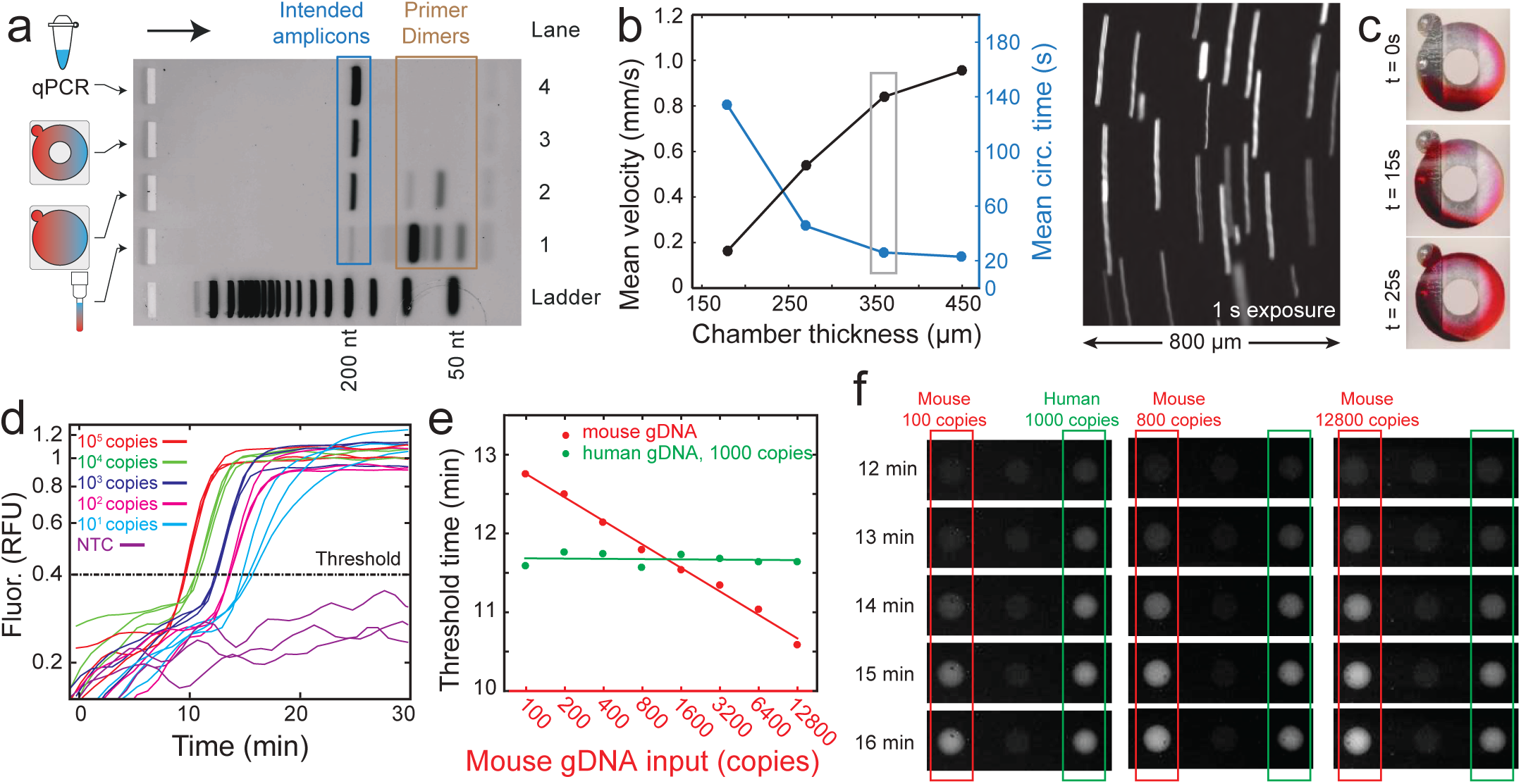
Donut PCR’s amplification specificity, sensitivity, and dynamic/quantitation range. **(a)** Donut PCR achieves dramatically higher PCR specificity than alternative implementations of convection PCR, enabling diagnostics-grade DNA analysis. Shown here are 2% agarose gel electrophoresis on single-plex PCR amplification products (human *TFRC* gene) using different platforms. Lane 1 shows results of convection PCR in a capillary tube using POCKIT instrument (GeneReach, Taiwan), based on principles described in ref. [22]. Hardly any intended amplicon molecules are generated, and the vast majority of amplification products are nonspecific amplicons or primer dimers. Lane 2 shows the results on the Donut PCR platform using a chip without the “donut hole.” There is significant nonspecific amplification and primer dimer formation due to the dead space in the middle with low temperature and circulation velocity. Lane 3 shows the results from the Donut PCR platform and chip. Lane 4 shows the results using a commercial qPCR instrument (Bio-Rad CFX96) in a 200 *μ*L PCR tube. **(b)** Dependence of circulation velocity on chamber thickness. Because the reaction solution undergoes laminar flow within the chamber, thicker chambers result in faster circulation. The right image is taken with 1 s exposure time and used for quantitation circulation velocity in 360 *μ*m thick chamber. The streaks show movement of fluorescent polystyrene beads (10 *μ*m in diameter). The variation in bead velocities likely correspond to the position in the z-axis, with beads closer to surfaces expected to be slower. **(c)** Pictures of the Donut PCR chip (360 *μ*m thick chamber) loaded with red dye to visualize fluid circulation. **(d)** Donut PCR sensitivity and dynamic range. As with qPCR, the observed Donut PCR spot fluorescence follows a sigmoidal curve. Here, we show fluorescence traces for amplification of human genomic DNA from 20 pg (10 copies) to 200 ng (10^5^ copies). The time at which the fluorescence crosses a threshold value (*T*_*t*_) is dependent on the log of the input DNA concentration, as expected. **(e)** Simultaneous amplification and quantitation of mixed samples of human and mouse genomic DNA. We designed primers and probes for the human *TFRC* gene and the mouse *Nono* gene, and quantitated each gene for samples wherein the human DNA was fixed at 1,000 copies and mouse DNA varied from 100 to 12,800 copies. The *T*_*t*_ values for human *TFRC* were unaffected by the changes in the mouse DNA concentration. See Supplementary Section S6 for additional data. **(f)** Sample fluorescence images of experiments shown in panel (e). The mouse probe (left spot) became brighter at earlier times when a larger amount of mouse DNA was used as input.

Next, we aimed to improve the amplification speed within the Donut PCR chamber because rapid turnaround is highly desirable for point-of-care applications such as pathogen identification. The speed of fluid circulation in the Donut PCR is impacted by the thickness of the reaction chamber, because in laminar flow, the fluid velocity near a surface is close to zero. Fig. 2b shows that mean circulation velocity increases in thicker chambers, consistent with expectations, and plateauing at roughly 360 *μ*m chamber thickness. At this thickness, for our standard chamber dimensions (10 mm outer diameter, 4 mm inner diameter), the chamber volume is approximately 25 *μ*L, consistent with commercial qPCR reaction volumes. With a 25 s circulation time, a standard 40-cycle PCR protocol can be completed within 20 minutes.

To evaluate the speed of PCR amplification and the dynamic range of the Donut PCR platform, we next ran the Donut PCR chip using different quantities of the NA18537 human cell line genomic DNA (Fig. 2d), ranging from 10 haploid genomic copies (5 cell equivalents, 20 pg) to 10^5^ haploid genomic copies. Triplicate repeat experiments showed consistent transition time (*T*_*t*_), defined as the time at which fluorescence first exceeds the threshold of 0.4 RFU. Furthermore, the *T*_*t*_ values for lower input DNA quantities increased as predicted in a log-linear fashion, similar to qPCR and demonstrating a dynamic range of at least 4 logs. All three negative control samples (water) did not have spot fluorescence exceeding the threshold, confirming specific PCR amplification and probe detection.

Next, we performed simultaneous quantitation of mouse and human DNA on a Donut PCR chip, in order to characterize the potential interference of quantitation accuracy due to presence of variable quantities of background DNA (Fig. 2ef). We observe that as the stoichiometric ratio of mouse DNA to human DNA ranges from 12.8 to 0.1; the value of *T*_*t*_ for human DNA is essentially unaffected. Simultaneously, the log-linearity of the mouse DNA *T*_*t*_ value is also unaffected by the presence of human DNA. Collectively, these results suggest that the Donut PCR platform could be used for multiplexed gene expression profiling.

### Spot position independence

The Donut PCR platform leverages the embedded pre-quenched microarray to achieve massive multiplexing. For these multiplexing capabilities to be realized in a diagnostic setting, it is necessary that the observed results are reproducible and consistent across different spots. To characterize inter-spot consistency, we constructed a 48-spot Donut PCR chip that includes quintuplet repeat spots for probes against 3 human genes, 3 rat genes, and 3 mouse genes, plus 2 positive controls and 1 negative control (Fig. 3a). We observe that although there is significant difference in endpoint spot fluorescence intensity, suspected to be primarily due to non-uniform illumination, the values of *T*_*t*_ are well conserved across all 5 replicate spots for each of the 9 genes.

**FIG. 3:**
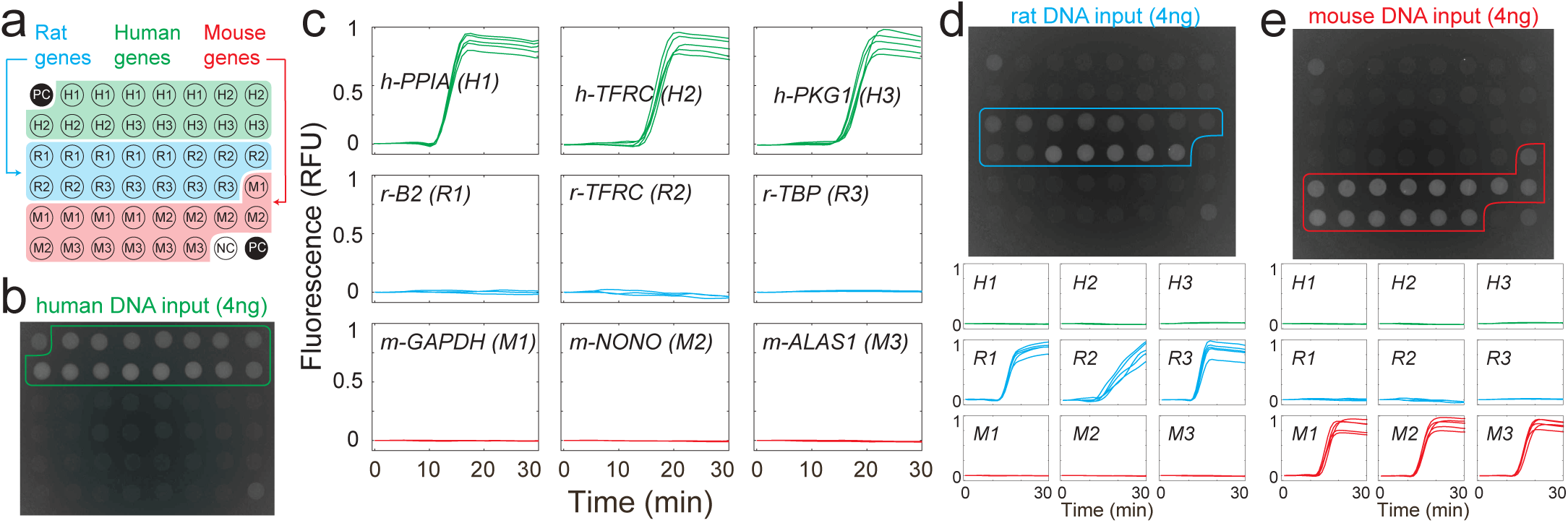
Spot position independence, established through redundant real-time detection of multiple genes. **(a)** Chip layout. Out of 48 spots, 3 were used as positive and negative controls (PC and NC), 15 spots were used for 5-fold replicate detection of 3 human genes (H1, H2, and H3), 15 spots for 3 rat genes (R1, R2, and R3), and 15 spots for 3 mouse genes (M1, M2, and M3). **(b)** Sample fluorescence image at t = 30 min when 4 ng of human gDNA was loaded as input, using PCR primers for all 9 genes (including 3 rat and 3 mouse). Only the human and positive control spots were bright, as expected. There is variation in the endpoint brightness of different probes due to both uneven illumination and due to differences in fluorophore coupling and probe spotting efficiency. **(c)** Time-course fluorescence traces for all 9 sets of probe spots. Threshold times are generally consistent across the 5-fold replicate spots. **(d)** Fluorescence image and time-course fluorescence using 4 ng rat DNA input, using the same primers and probes as in panels (b) and (c). **(e)** Fluorescence image and time-course fluorescence using 4 ng mouse DNA input.

### Portable instrument

Experiments presented thus far have used a commercial fluorescence microscope as the readout instrument. To facilitate the adoption of the Donut PCR platform for a variety of applications that require portability and rapid turnaround, we next designed and built a portable Donut PCR instrument (Fig. 4a). Importantly, commercial qPCR instruments require wall power, preventing them from being rapidly deployed to point-of-care or field-use settings where rapid diagnostic testing may be needed. A major component of the power requirement is the need to rapidly cool PCR reaction mixtures from 95 °C (denaturing step) to 60 °C (annealing step). Because Donut PCR achieves rapid thermal cycling using passive fluidic flow, it requires only constant temperature heaters with low power consumption. Our breadboard prototype (Fig. 4b) thus is able to run off a mid-size 12V battery (12V × 18 Ah) for 9-10 experiments, and does not require connection to an external power source.

**FIG. 4:**
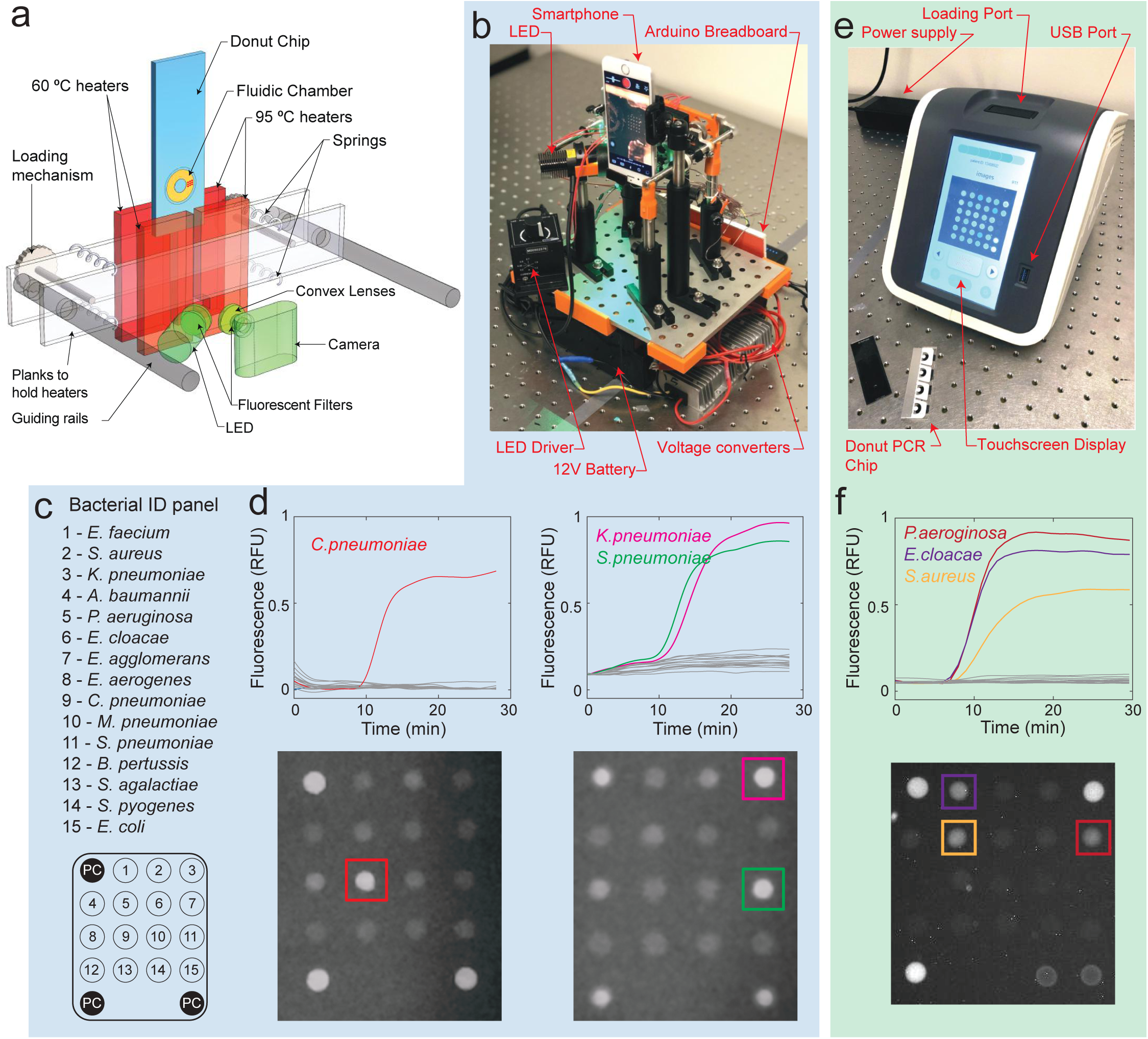
Donut PCR instruments and bacterial identification panel. **(a)** Simplified schematic of the loading, heater-clamping, and imaging modules. **(b)** Picture of initial breadboard prototype instrument, using an iPhone 6 as a portable camera. This prototype is fully battery-powered (12V) and does not require an external power supply. **(c)** Probe layout for a Donut PCR chip designed to detect bacterial species frequently observed in nosocomial (hospital-acquired) infections. **(d)** Experimental results on the breadboard prototype using a 0.5 ng clinical isolate sample of *C. pneumoniae* (left) and a mixture of clinical isolate samples of 0.25 ng *K. pneumoniae* and 0.25 ng *S. pneumoniae* (right). **(e)** Picture of functional prototype instrument. **(f)** Experimental results on the functional prototype using a mixture of clinical isolate samples consisting of 1 ng *P. aeroginosa*, 1 ng *E. claocae*, and 1 ng *S. aureus*. For historical reasons, input quantity and chip layout differed from panels (c), and (d). See Supplementary Section S9 for probe layout.

There are 5 main modules in the instrument: (1) a mechanical mechanism to mount the chip against the heaters with sufficient force to ensure good thermal contact, (2) a closed-loop thermal system to ensure the chip is differentially heated to the desired temperatures, (3) an optics setup to illuminate the chip and filter out scattered light to reduce fluorescence background, (4) a camera to acquire fluorescence images, and (5) a microcontroller to coordinate timing of all components. In total, the cost of the components of this breadboard prototype was roughly $2,500, with the bulk of the cost from to the camera (an iPhone 6S, $600), the optical filters (ThorLabs MF530-43 and Chroma AT575LP, $500), LED light source (ThorLabs LED1B, $325), and LED driver (ThorLabs M530L4, $296).

To validate the functionality of this prototype instrument, we constructed a 15-plex bacterial identification panel on the Donut PCR chip. The DNA sequence encoding the 16S ribosomal RNA and 23S ribosomal RNA in bacteria are mostly conserved but contains 9 hyper-variable regions with sequences that differ across species but are conserved within strains of the same species. For this 15-plex bacterial panel, we constructed 15 different probes that target distinct 16S sequences (located in the V3 hyper-variable region) that serve as signatures for 15 important bacterial species frequently implicated in nosocomial (hospital-acquired) infections, including the most common ESKAPE [23] bacteria. Nosocomial infections cause roughly 20,000 deaths in the United States per year [24], and rapid pathogen identification could help inform timely targeted antibiotic treatment that can improve patient outcomes, limit spread, and drug resistance [17, 18].

Our ESKAPE panel performed as we expected on the bread-board prototype instrument, successfully identifying both a single bacteria species and a combination of two bacteria species. Because the current chip and instrument design does not include sample preparation and DNA extraction modules, we performed validation using DNA input from clinically derived isolates and reference strains purchased commercially from ATCC collection and Zeptometrix Corp. (Buffalo, NY). We individually tested each DNA sample obtained (see Supplementary Section S9 for additional experimental data on the 15-plex bacterial panel using the breadboard prototype).

Because the breadboard prototype is open to the air, it is vulnerable to ambient dust that can occlude or distort the optics and ambient light that reduces image signal to noise ratio. We next contracted a third-party engineering firm to build a closed instrument with similar functionality to our breadboard prototype (Fig. 4e). As expected, the signal to noise ratio of the fluorescent spots were improved, and the 15-plex bacterial identification panel was able to clearly identify all 3 bacterial species in a mixed sample of clinical isolates (Fig. 4f). See Supplementary Section S9 for revised chip layout for the production prototype instrument.

### Single nucleotide polymorphism (SNP) genotyping

Single nucleotide polymorphisms are natural variations in the human genome; over 100 million distinct SNP loci have been reported in the human genome [25, 26]. Genome-wide association studies (GWAS) have discovered a wide range of SNPs that correlate with disease risk, from diabetes [27], neurodegnerative disease [28, 29], hereditary breast [30] and colorectal cancers [31], and coronary disease [32]. In addition to human disease-related applications, SNP genotyping can also be used for DNA forensics application [33] as well as agricultural seed selection [34].

Currently, the microarray are the preferred technology for SNP genotyping applications, due to its massive multiplexing, high automation, and acceptable economics (typically ≤$60 per Affymetrix array chip). However, analyzing a sample using a microarray takes approximately 24 hours. Furthermore, because it requires 3 large specialized instruments for hybridization, washing, and imaging, samples need to be transported to central labs for analysis, adding more days to the total turnaround time. While a 3-7 day turnaround is acceptable for non-time-sensitive applications, other SNP genotyping applications require rapid turnaround. One prominent example is pharmacogenetics [15]; the use of SNP genotyping information to inform dosage of drugs such as warfarin based on individualized drug metabolism rates [35]. Information to guide accurate dosage of warfarin to treat patients suffering from stroke or deep vein thrombosis needs to be provided immediately.

For SNP genotyping, we use the X-probe architecture [20] to achieve probe-based single-nucleotide discrimination while limiting the reagent costs of chemically modified DNA (Fig. 5a, see also Supplementary Section S10). For each SNP locus, we design two separate surface-bound X-probes, one to each allele. Human DNA samples that are homozygous at the SNP locus will only have the corresponding allele spot light up, while samples that are heterozygous will have both spots light up (Fig. 5b). The overall workflow from buccal (cheek) swab sample collection takes less than 1 hour, including less than 5 minutes of hands-on time (Fig. 5c). See http://demovideo.torus.bio for a 2-minute video of the entire workflow.

**FIG. 5:**
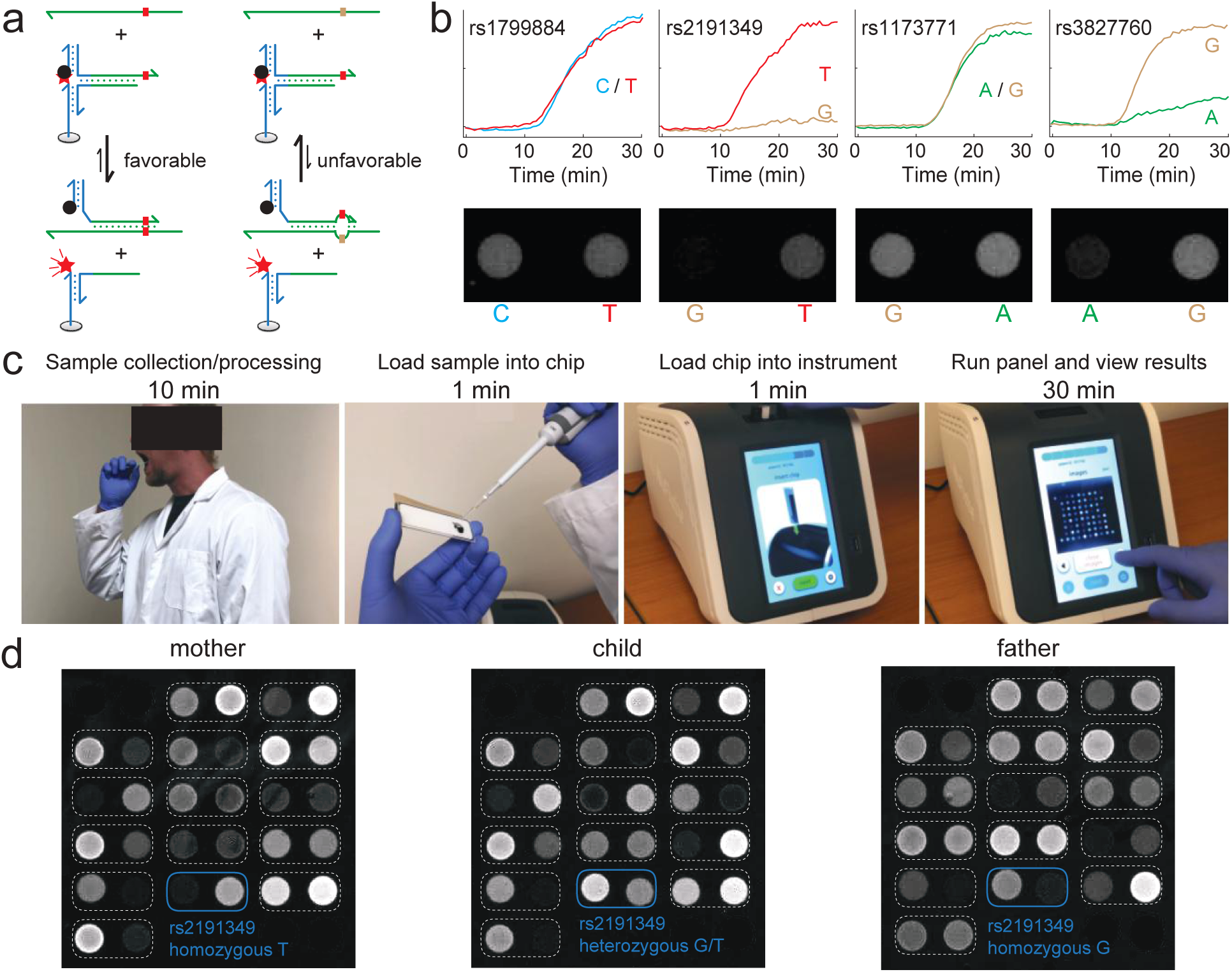
Single nucleotide polymorphism (SNP) genotyping using Donut PCR. **(a)** To simultaneously achieve single-nucleotide discrimination and reduce functionalized oligo costs, we use the X-probe architecture [20] for the SNP genotyping panel here. Due to the thermodynamics and kinetics of the X-probe design, it will bind only perfectly matched DNA sequences, and even DNA sequences differing by a single SNP allele will not favorably bind to the probe. **(b)** Sample experimental results on 4 SNP loci. Each pair of probes performs genotyping on a different SNP. **(c)** Illustration of workflow for human buccal swab sample testing on the Donut PCR. Total turnaround time is under 1 hour. **(d)** Genotyping 15 SNP loci from buccal swab samples from family trio. See Supplementary Section S10 for SNP loci covered, probe layout, time-based fluorescence plots. Buccal swab and finger-stick samples were collected with informed consent.

To showcase the validation and accuracy of SNP genotyping by the Donut PCR platform, we constructed a 30-spot array corresponding to the alternate alleles for 15 different human SNP loci. Initial panel testing showed correct SNP genotype calls for all 15 loci for 3 different human cell line gDNA samples. Next, this panel was applied to buccal swab samples from a family trio of mother, father, and child under informed consent, using the production prototype instrument shown in Fig. 4e. Endpoint fluorescence images of the samples are shown in Fig. 5d, but as usual SNP genotype calls are made using time-based fluorescence traces.

## Discussion

The Donut PCR platform presented here achieves rapid, sensitive, and quantitative detection of many DNA targets from a single sample using a closed, portable, and affordable instrument. Although we have limited our studies in this work to arrays of 16 to 48 spots, probe printing is a highly scalable process and we have demonstrated that over 1000 probes can be printed on a single Donut PCR chip (Supplementary Section S11). We believe that the Donut PCR platform will be competitive in a range of applications where rapid and highly multiplexed DNA detection is needed in decentralized settings.

With the recent coronavirus pandemic, one apt use case for the Donut PCR may be in disease surveillance, e.g. at airports. RNA viruses such as Covid-19, influenza, and HIV are particularly prone to genetic drift due to the high error rates of reverse transcriptase, and the NextStrain database reports thousands of genetic variants for each virus [36]. Consequently, single-plex isothermal amplification assays [9, 10] are vulnerable to low and decreasing clinical sensitivity as virus genomes evolve away from the target sequences singleplex assays are designed to detect. The high multiplexing of the Donut PCR platform can overcome potential clinical false negatives by redundantly detecting many conserved pathogen-specific RNA or cDNA sequences.

The single nucleotide specificity demonstrated by the Donut PCR platform in Fig. 5 allows simultaneous detection of antibiotic resistance with pathogen identification. Antibiotic resistances are typically caused either by gain of a gene (e.g. mecA for methicillin resistance) or mutation of a gene (e.g. gyrA for fluoroquinone resistance [37]). While other multiplex PCR platforms such as BioFire [11] or StatDx [38] can detect drug resistance due to gain of gene, they typically lack the molecular specificity needed to identify resistance caused by point mutations. The ability to rapidly identify infections and prescribe effective antimicrobial therapies is especially needed to reduce the mortality and morbidity rate of nosocomial infections in the United States.

The core innovations in the Donut PCR platform are the donut-shaped chamber to allow reliable and low-power convection-based qPCR, and the integrated pre-quenched microarray to allow massively multiplexed readout through spatial separation. As a result, our experimental demonstrations in this manuscript were “DNA in, answer out” workflows. We did not consider sample preparation and DNA extraction modules. Extensive prior work have reported a variety of different lab-on-a-chip approaches to process blood, nasal/buccal swab, urine, and cerebrospinal fluid. Integration with a pre-analytical module will likely be necessary for adoption of the Donut PCR platform in point-of-care settings for infectious disease diagnostic applications.

Another potential application area for Donut PCR is multigene expression analysis. Cancer prognosis tests such as Oncotype Dx [39] use tumor gene expression profiles to stratify patients for more aggressive treatment such as chemotherapy. Recently, significant research on host biomarkers [40, 41] also suggests that human gene expression can be used to differentiate exposure to bacteria vs. viruses, and could serve as a complementary set of markers to pathogen DNA. We believe that Donut PCR’s ability to rapidly and simultaneously detect and quantitate many different nucleic acid markers positions it well as a platform for performing complex DNA and RNA diagnostics in settings convenient to the patient.

## Acknowledgements

This work was supported by NIH grant R01CA203964 to DYZ. The authors thank Jianyi Nie for editorial assistance. The authors thank David Walt for suggestions on instrumentation design. The authors thank Torus Biosystems for lending production prototype instruments used to collect data in Fig. 4ef and Fig. 5.

## Author contributions

DK and DYZ conceived the project. DK and DYZ performed primer and probe design. DK performed chip design and construction. DK, JL, and JXZ performed instrument design and construction. DK and DYZ wrote the manuscript with input from all authors.

## Additional information

We have complied with all relevant ethical regulations. Correspondence may be addressed to DYZ. There are patents issued on toehold probes and X-probes used in this work. There are patents pending on the Donut PCR chip and Donut PCR instrument presented in this work. DK, JL, and DYZ declare competing interests in the form of employment (DK) or consulting (JL and DYZ) for Torus Biosystems. JXZ and DYZ declare competing interests in the form of employment (JXZ) or consulting (DYZ) for Nuprobe. DYZ declares a competing interest in the form of consulting for Avenge Bio.

## Software and Data Availability

Raw fluorescence images/movies and code for image analysis are available upon request.

