## Supplementary info for "Donut PCR: a rapid, portable, multiplexed, and quantitative DNA detection platform with single-nucleotide specificity" (from another version of this preprint: version 2, 2020-05-02)

**Contents**

**S0. Material and Methods** 2
Surface functionalization 2
DNA oligonucleotides 3
Design of primers 3
Reference DNA samples 3
DNA extraction 3
Agarose gel electrophoresis 3
Microarray printing 3
General considerations on thermodynamics of toehold and X-probes 4
Composition of PCR reaction 4

**S1. Assembly of Donut PCR fluidic chamber** 5

**S2. Design and structure of the research-grade real-time Donut PCR thermocycler** 6
Optical module 6
Thermal module 6
Mechanical module 6

**S3. Image processing and quantitation** 10

**S4. Polymerase selection** 14

**S5. Fluid dynamics in Donut PCR chamber** 15
Distribution of flow velocities in the convection flow 15
Flow geometry in three different convection flow chambers 15
Donut PCR amplification speed depends on chamber thickness 16

**S6. Quantitation of DNA targets based on $T_t$** 16

**S7. Simultaneous amplification and detection of 20 genetic loci from human genomic DNA** 18

**S8. Battery-Powered Breadboard Donut PCR instrument** 21

**S9. Bacterial identification panel** 22
Design consideration for 16S-based bacterial identification 22
Detection of 6 bacterial species' DNA using battery-powered Donut PCR instrument 23
Detection of 15 bacterial species' DNA using functional prototype Donut PCR instrument 24

**S10. Single nucleotide polymorphism (SNP) genotyping.** 27
SNP genotyping using buccal swabs 29

**S11. Donut PCR capabilities for further development.** 30
Ultrafast Donut PCR 30
Feasibility of scaling to massively multiplexed panels 31

**References** 31

### S0. MATERIAL AND METHODS

#### Surface functionalization

Attachment of the pre-annealed probe to the surface of the fluidic chamber is performed via strain-promoted azide-alkyne cycloaddition (SPAAC) between dibenzocyclooctyne (DBCO) functionalized probe and azide ($N_3$) functionalized surface of the fluidic chamber. $N3$ surface functionalization is performed according to the following protocol depicted in Fig. S1a:

- Step (i). Glass slides are treated with sodium hydroxide solution (10 M) for 30 min in Branson ultrasonic bath (Bransonic CPXH Digital Bath 2800) at the high power of ultrasound generation;
- Step (ii). After washing with copious amount of water and drying at 120 °C, the slides are treated with oxygen plasma for 30 min in Harrick plasma cleaner at high power and maintaining oxygen pressure in range of 0.2 - 0.3 torr;
- Step (iii). Hydroxylated slides are treated with a 5% solution of N-(2-aminoethyl)-2,2,4-trimethyl-1-azasilacyclopentane (Gelest) in dry dichloromethane (Sigma) for 16 hrs at room temperature, followed by washing with 95% ethanol in water solution; amino-salinized slides are baked in ambient atmosphere at 105 °C for 2 hrs;
- Step (iv). Functionalization of the amino-terminated slides with isothiocyanate (ITC) moiety is performed with 10 mM solution of *p*-phenylene diisothiocyanate (Sigma) in dry pyridine-dimethylformamide (both chemicals are from Sigma) mixture (9:1, v:v) for 2 hrs at room temperature, followed by washing with 95% ethanol in water solution and drying at 80 °C for 1 hr ;
- Step (v). Azido-functionalization of the ITC slides is performed at 37 °C for 2 hrs with a solution of 11-azido-3,6,9-trioxaundecan-1-amine (50 $\mu$L) in 0.1 M phosphate buffer pH 8 (10 mL), followed by washing with copious amount of water and drying 45 °C.

At this stage the slides are ready for microarray printing or can be stored vacuum-sealed at 4 °C for at least 6 months without loss of conjugation activity.

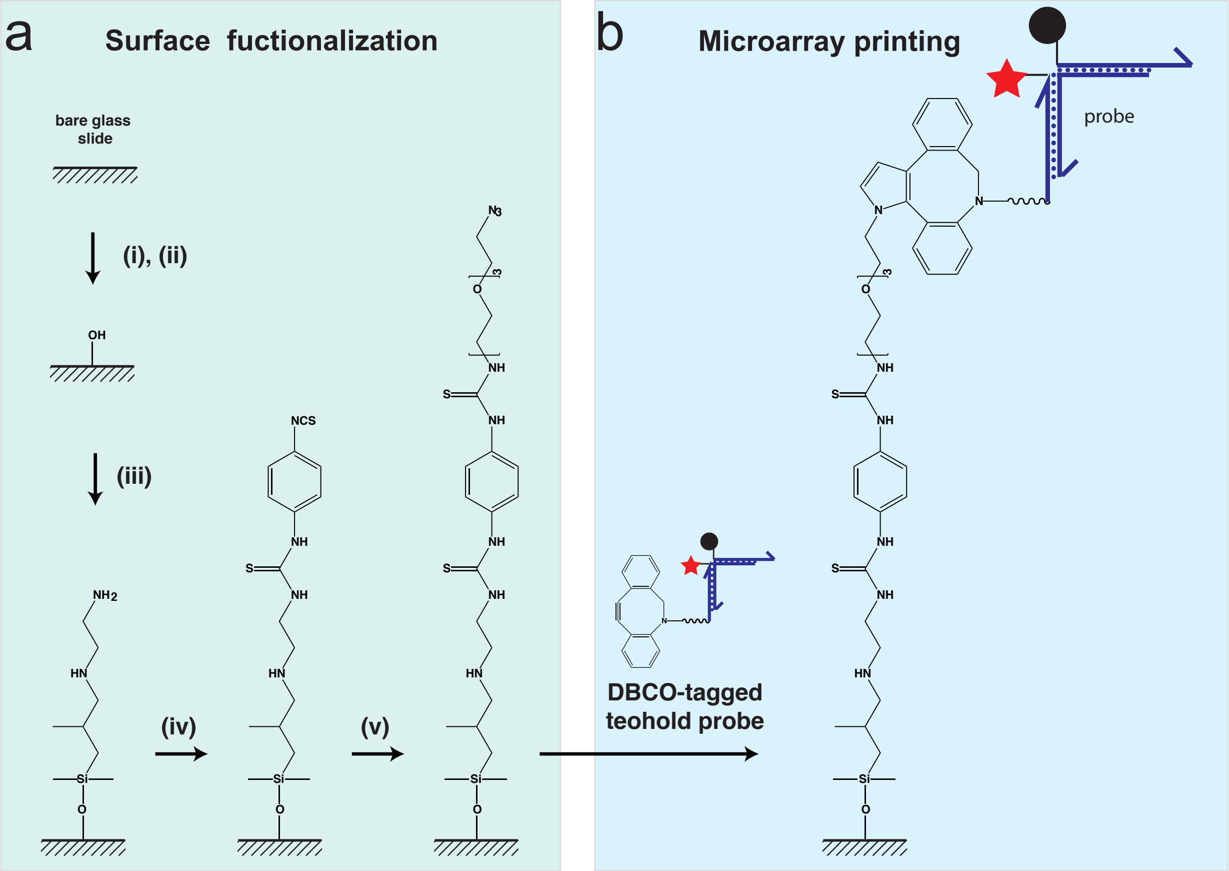


FIG. S1: Pipeline for surface functionalization and microarray printing **(a)** Surface functionalization includes four main stages. First, bare class slides (soda-lime) are (i) treated with NaOH (ii) and exposed to oxygen plasma. This allows the creation of a highly hydrophilic surface with a large amount of silanol groups. Next, the surface is (iii) converted into an amine functionalized surface by reacting with N-(2-aminoethyl)-2,2,4-trimethyl-1-azasilacyclopentane. Then, step (iv) introduces amino reactive moiety via reaction with *p*-phenylene diisothiocyanate. Finally, the azido group ($N_3$) is introduced by reacting with 11-azido-3,6,9-trioxaundecan-1-amine (v). **(b)** DBCO-functionalized pre-annealed probes are printed using non-contact printing method. Functionalization of the probes to the surface is governed by strain promoted azide-alkyne cycloaddition (SPAAC) mechanism. Importantly, the deactivation of the unreacted $N_3$ groups is not required, since $N_3$ group has no bioorthogonal "partners" in complex biological environments.

### DNA oligonucleotides

All DNA oligonucleotides used in the study were purchased from Integrated DNA Technology (Coralville, IA), except for the strands labeled with BHQ-2 moiety which were purchased from Biosearch Technologies (Petaluma, CA). All PCR primers were ordered without purification, but desalted. All fluorophore-labeled oligonucleotides, quencher-labeled oligonucleotides, and unlabeled oligonucleotides shorter than 50 nt were ordered with reverse-phase HPLC purification. All unlabeled oligonucleotides longer than 50 nt were ordered with PAGE purification.

### Design of primers

Main considerations during primer design were as follows: 1) amplicon length was restricted to 300 bp or shorter; 2) standard free energy of primer binding ($\Delta G^{\circ}$) was in the range of -11 to -12 kcal/mol at 60 °C at 60 °C in a solution with 6mM $Mg^{2+}$ concentration; 3) no predicted intermolecular binding with standard free energy more negative than -3 kcal/mol; 4) no intramolecular secondary structures with standard free energy more negative than -5 kcal $mol^{-1}$.

### Reference DNA samples

Reference human genomic DNA samples were purchased from Coriell Institute for Medical Research, Camden, NJ
Reference bacterial genomic DNA samples were purchased from ATCC collection, Gaithersburg, MD, and Zeptometrix Corporation, Buffalo, NY

### DNA extraction

DNA extraction from fresh fingerstick blood samples (200 $\mu$L) was performed using Qiagen blood mini kit strictly following manufacturer's protocol. DNA extraction from buccal swab samples was performed using QuickExtract DNA Extraction Solution from Lucigen (recently acquired by Biosearch Technologies). In brief: after collecting buccal swab samples (SK-2S buccal swab, Isohelix, Kent, UK), the swabs were placed and vortexed in 0.5 ml of QuickExtract DNA Extraction Solution. The solution (with the swabs) was heated at 60 °C for 10-15 min, at 95 °C for 3 min, and then cooled down to room temperature. Two micro-liters of the solution was used as DNA input for SNP genotyping assays.

### Agarose gel electrophoresis

Agarose electrophoresis was performed in 1×TEA buffer (40 mM Tris-HCl, 20 mM Acetic acid and 1 mM EDTA, pH 8.6) at the electric field strength of about 10 V/cm.

### Microarray printing

**Probe annealing.** Toehold microarray probes were annealed in 2×PBS buffer according to the following protocol:
90 °C for 3 min, then uniformly decreasing temperature down to 30 °C over a period of 1 hr. Concentrations of the probe components were as follows:

Universal Probe Architecture (Fig. S2a):
- Anchor strand - 2 $\mu$M
- Arm strand - 3 $\mu$M
- Quencher strand - 4.5 $\mu$M.

Universal Probe Architecture (Fig. S2b):
- Anchor strand - 2 $\mu$M
- Arm strand - 3 $\mu$M
- Quencher Arm strand - 4.5 $\mu$M.
- Universal Quencher strand - 8 $\mu$M.

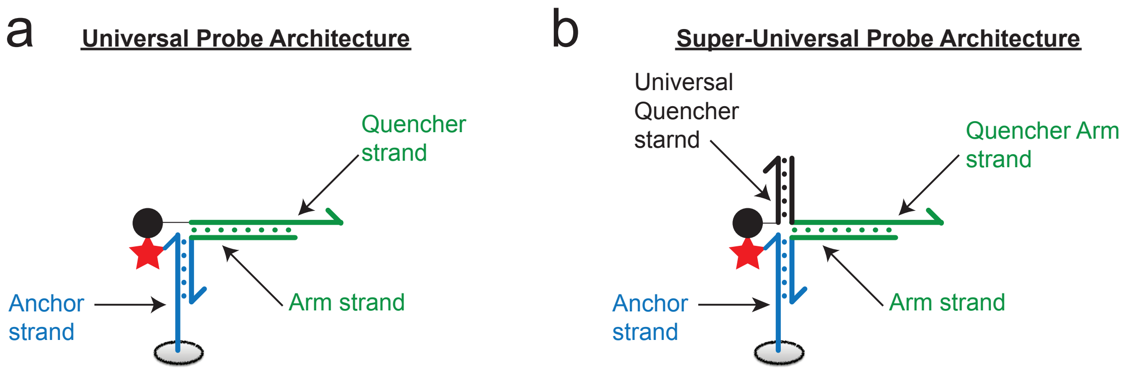


FIG. S2: DNA architecture of the **(a)** toehold probes and **(b)** X-probes used in the study. Both probe architectures comprised a universal Anchor strand for tethering to the surface via DBCO moiety and was tagged with a fluorescent dye (AttoTech-550) at its 3' end, which served as a fluorescence reporter. The Anchor strand was designed to be universal to allow hybridization with different Arm strands bearing 3' domain complementary to the Anchor. The Arm and the Quencher strands were target specific and responsible for target recognition and SNV discrimination. For the super-universal probe, the Quencher arm strand contained a 5' domain (colored in black) used for hybridization with a universal quencher oligonucleotide, while the universal probe architecture possesses specific quencher arm labeled with fluorescent quencher. Both probe architectures allowed us to significantly reduce costs and time for assay reconfiguration by avoiding expensive and long-lead-time dual labeled oligonucleotides.

**Microarray spotting solution preparation**. Microarray spotting solution was prepared by diluting the annealed probes with Tris-EDTA buffer (TE) spiked with $MgCl_2$ to obtain the following final concentration of the components:
- 1×TE buffer (10 mM Tris, 1 mM EDTA)
- 10 mM $MgCl_2$
- 0.5 $\mu$M of the probe, recalculated to the concentration of the Anchor strand.

**Microarray printing**. Microarrays were printed using Scienion S3 microarray printer equipped with PDC070 micro-nozzles. Printing was preformed at 70% relative humidity maintained in the printing enclosure. After the printing run finished, the arrays were allowed to rest for 15 minutes at the same humidity and were then dried in a desiccator under reduced pressure for 1 hr. The microarrays were stored at room temperature in the vacuum-sealed polyethylene bags until final assembly and use.

### General considerations on thermodynamics of toehold and X-probes

During the probe design we kept the same toehold binding energy within a range -9.5 to -10.5 kcal mol$^{-1}$ (for all probes designs) for a fully complementary sequence. By comparison, toehold binding with variant amplicon sequence would have higher energy, thus not favorable for toehold displacement reaction initiation, which in turn would result in no or weak fluorescence signal increase. The binding energetics for the specific arm sequence (binding between the Arm strand and the Quencher strand or Quencher Arm strand) was kept in the range of -22 to -24 kcal mol$^{-1}$ to prevent spontaneous dissociation of the quencher strand from the probe. Both Anchor strand and Universal quencher strand had binding energy of around -24 kcal mol$^{-1}$. Probes designed for SNP discrimination were also equipped with a unique non-homologous double-stranded region with binding energy of around - 5 kcal mol$^{-1}$. Presented probe energetics was calculated for 60 °C and concentration of $Mg^{2+}$ cations at 6 mM, using the script written in our laboratory.

### Composition of PCR reaction

**Donut PCR** was performed using 15 Units of AptaTaq Δ exo DNA Polymerase (Roche) per reaction, 1×AptaTaq Fast PCR buffer and final $MgCl_2$ concentration of 6 mM. The increased concentration of magnesium ions was reasoned by their nonspecific absorption on glass surface. The concentration of each of the standard deoxynucleotide triphosphates (dATP, dTTP, dGTP, dCTP) was 0.25 mM. Each PCR reaction mixture was supplied with 5% DMSO and BSA at a concentration of 0.3 $\mu$g $\mu$L$^{-1}$. Concentrations of primers varied from 50 to 400 nM depending on the assay run. The Donut PCR instrument heaters were set to 60°C (array side) and 95 °C.

**Standard (tube) PCR** was performed using AptaTaq Δ exo DNA Polymerase (Roche) with 1×AptaTaq Fast PCR buffer at a final $MgCl_2$ concentration of 1.5 mM. The concentration of each of the standard deoxynucleotide triphosphates (dATP, dTTP, dGTP, dCTP) was 0.2 mM. Concentration of primers was 200 nM. PCR amplification regime of 95°C for 1 min, 30 cycles of 95°C for 10 s, 60°C for 10 s, 72°C for 15 s, and a final elongation of 72°C for 30 s was used.

### S1. ASSEMBLY OF DONUT PCR FLUIDIC CHAMBER

A complete Donut PCR fluidic chip assembly pipeline is shown in Fig. S3. After microarray was printed, it was assembled with a cover slide and a polymer spacer. Cover slide was a standard 1 mm thick microscope slide with two drilled holes. Drilling was performed with diamond micro-bites using Dremel 3000 variable rotary kit according to the drawings shown in Fig. S4a. After drilling, the slide was treated with 10 M NaOH solution for 30 min, washed with copious amount of water, and dried. Polymer spacer was made from layered 3M 9965 medical tape cut to the desired geometry using Universal Laser System X-600. Thickness of the single layered spacer was measured to be 90 $\mu$m. Thus, to fabricate Donut PCR cartridge with 180, 270, 360 or 450 $\mu$m thick chamber, - 2, 3, 4 or 5 layers of the polymer spacer, respectively, were laminated before laser cutting. Beside a one-chamber Donut PCR chip, we developed a Donut PCR Chip with 4 reaction chambers that could be used for running up to four reactions simultaneously (it was obvious that the microarray slide had to have four microarray clusters printed). This procedure could be performed on the research-grade Donut PCR thermocycler equipped with precisely controlled XY translation stage (discussed below). Drawing of the cover slide with loading ports and a polymer spacer are presented in Fig. S4b.

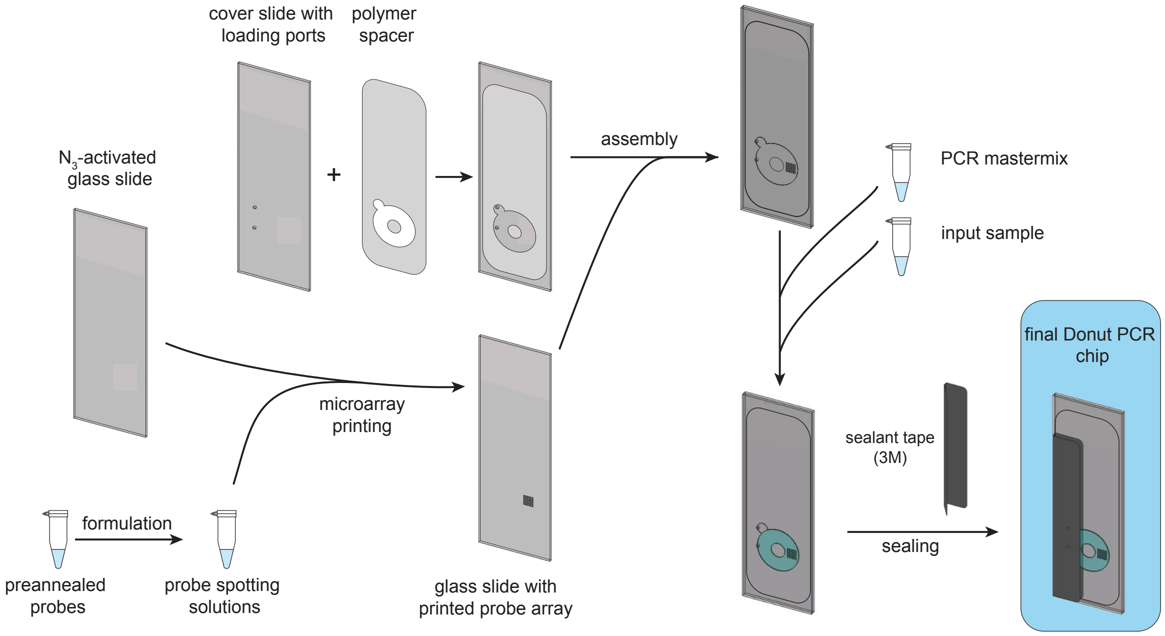


FIG. S3: Donut PCR fluidic chip with one reaction chamber assembly pipeline. The reaction chamber was made of two microscope glass slides separated by a 360 $\mu$m thick double-sided medical adhesive spacer (3M 9965 medical tape) with a circular opening. A circular insert made of the same adhesive tape was located in the middle of the opening and turned the chamber into a "donut-shaped" reservoir with a volume of 25 $\mu$L. The fluidic chamber also had two loading ports in one of the slides for PCR mixture loading and air bypass. Once the chip was assembled, it was loaded with PCR mastermix mixed with input DNA sample and sealed with 3M sealing tape.

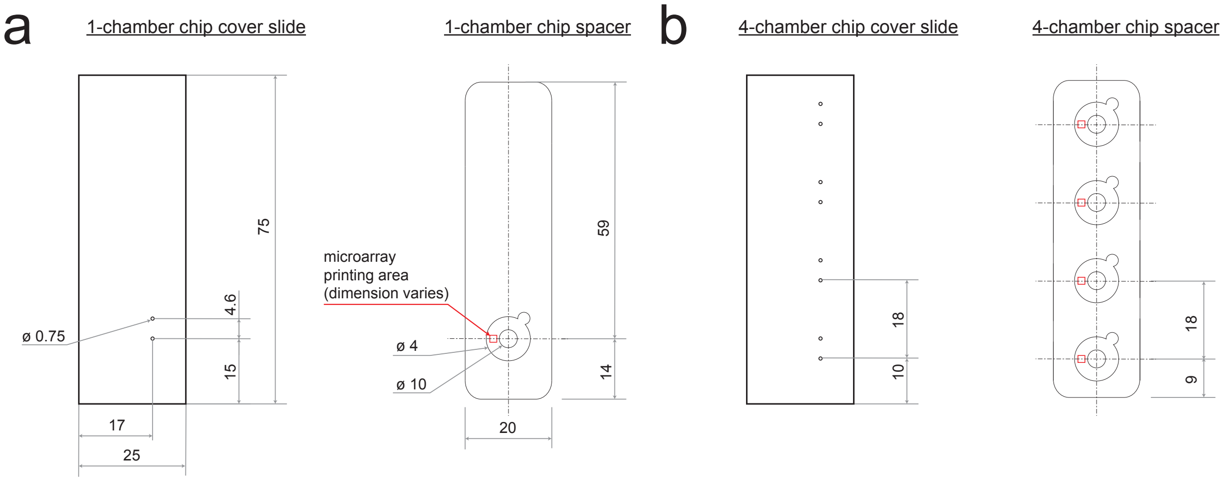


FIG. S4: Drawings of **(a)** 4-chamber and **(b)** 1-chamber Donut PCR chip cover slides and polymer spacers. All measurements presented are in mm.

### S2. DESIGN AND STRUCTURE OF THE RESEARCH-GRADE REAL-TIME DONUT PCR THERMOCYCLER

We designed and made a research-grade real-time Donut PCR thermocycler. The instrument is capable of maintaining desired temperatures and acquiring fluorescent images from one Donut PCR chip possessing multiple Donut fluidic chambers.The instrument comprises three main units: an optical module, a thermal module, and a mechanical module.

#### Optical module

The optical module of the instrument represents a modified Zeiss Axiovert M1 inverted epifluorescent microscope. In particular, the following modifications were made: (a) 30mm cage mounted dielectric mirror (ThorLabs Part CCM1-E02) was attached to the microscope turret via extension lens tube (ThorLabs Part SM1S25); (b) in order to prevent microarray elements from constant light illumination and, as a consequence, photobleaching, the caged mirror was equipped with an optical beam shutter (ThorLabs Part SH1/M); (c) finally, a genuine Zeiss 5x EC Plan-NEOFLUAR objective was connected to the shutter via a set of lens connectors. The executed modifications allowed imaging of the vertically mounted microarrays to have a field of view of 2.77 × 2.08 mm (Fig. S5ac).

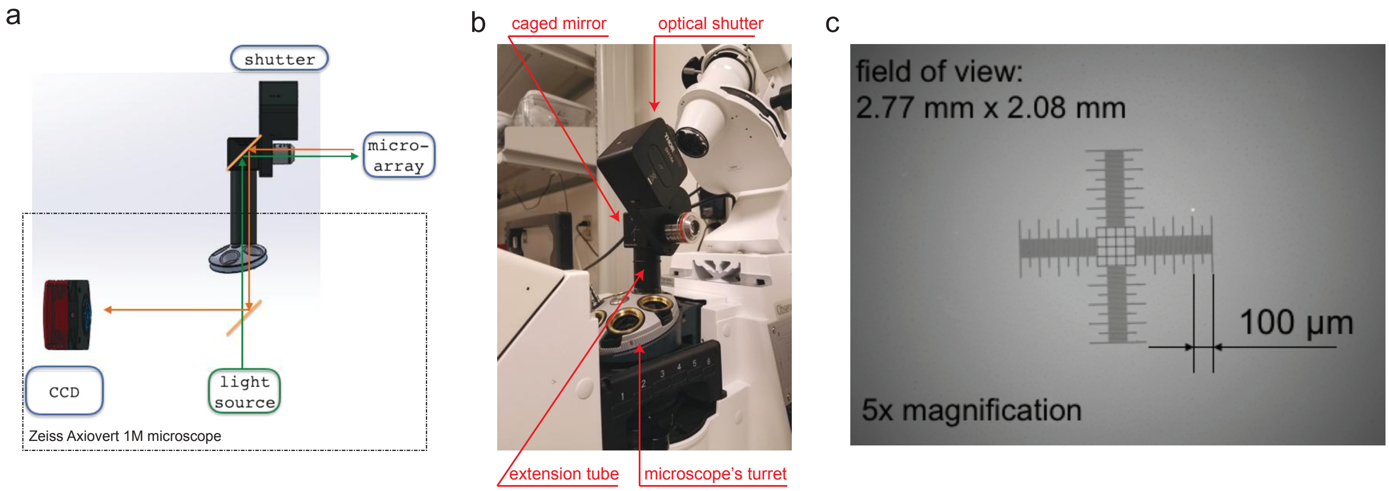


FIG. S5: Optical system of the research-grade Donut PCR instrument. **(a)** Schematic and light paths of the research-grade optical system. Zeiss microscope' fluorescent filters are intentionally not shown here to avoid overloading the drawings. **(b)** Photo of the modified Zeiss Axiovert M1 fluorescent microscope externally equipped with the extension tube, caged mirror and a solenoid based beam shutter (all parts from TorsLabs). **(c)** Bright-field photo of the size calibration slide.

#### Thermal module

The thermal module of the instrument is simple and is based on aluminum plates with glued Kapton foil heaters (Omegalux, KHVL-0504/10-P). For the research-grade real-time Donut PCR thermocycler, only two heaters provide required temperature settings as shown in Fig. S6a. The heaters are mounted on a heater holder (3D printed PLA) and has embedded temperature sensors (PerfectPrime, TL0225 K-Type thermocouple flat surface contact) connected to the PID temperature controllers (Inkbird, ITC-100VH). Both heaters are powered with 19 DCV power supplies via a solid state relay (Kyotto, KF0602D). Air fans shown in Fig. S6a are used only for cooling the heaters after the amplification reaction is finished and are not used to maintain preset temperatures. Accuracy of the design temperature regulating heating unit spans a range of $\pm 0.4$ °c. Fixation of the Donut PCR chip consumable on the heaters is performed using 2.5 cm wide paper clips, as shown in the Fig. S6b (in actual practice, six paper clips were used to provide better thermal contact between the chip and the heaters).

#### Mechanical module

The mechanical module of the instrument is designed to allow real-time imaging of multi-chamber Donut PCR chips by its aligning with the microscope's objective. As shown in Fig. S7 (drawings) and S8 (photos), the heating unit is held by the 3-D printed heating unit holder which, in turn, is fixed to the Z-axis motorized linear translation stage (ThorLab Part LTS150/M). The Z-stage is responsible for vertical alignment of the multi-chamber Donut chip. To allow horizontal alignment of the Donut chip, we mounted the Z-axis translator to the motorized X-axis linear translation stage (ThorLab

Part PT1/M-Z8). The travel ranges of Z- and X-linear translation are at 150 and 25 mm, respectively. Focusing of the Donut chip is realized using manual translation stage (Y-axis, ThorLab Part PT1A/M). Bringing Donut chip in the microscope's objective focus is usually performed at the very beginning of image acquisition cycle and does not require adjustment during the run.

Operation of optical shutter, Z-, and X- linear translators are controlled via corresponding driver controllers. Matlab scripts available upon the request.

a

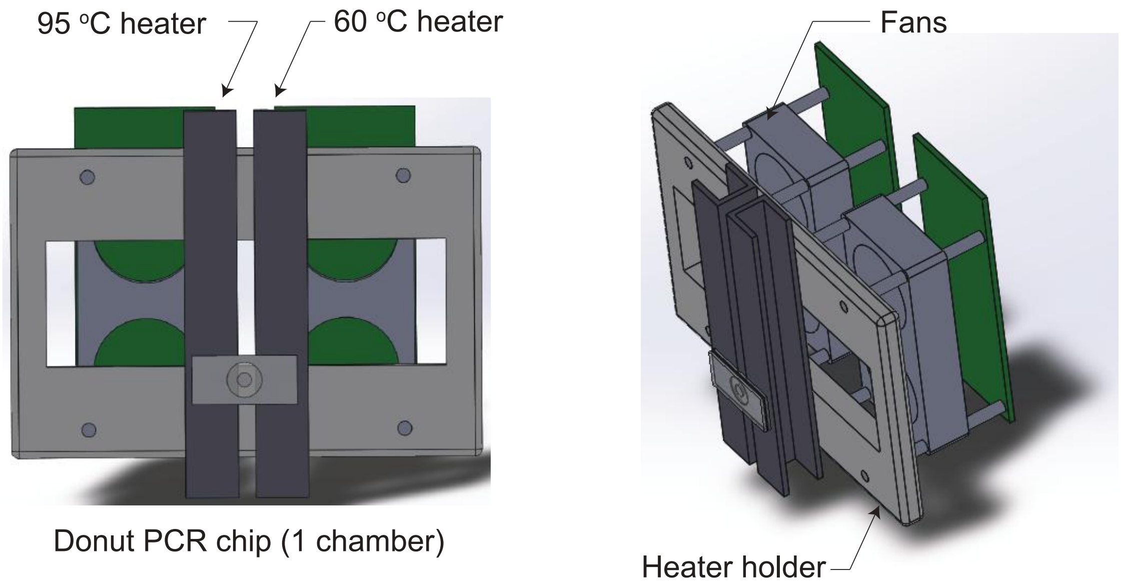


b

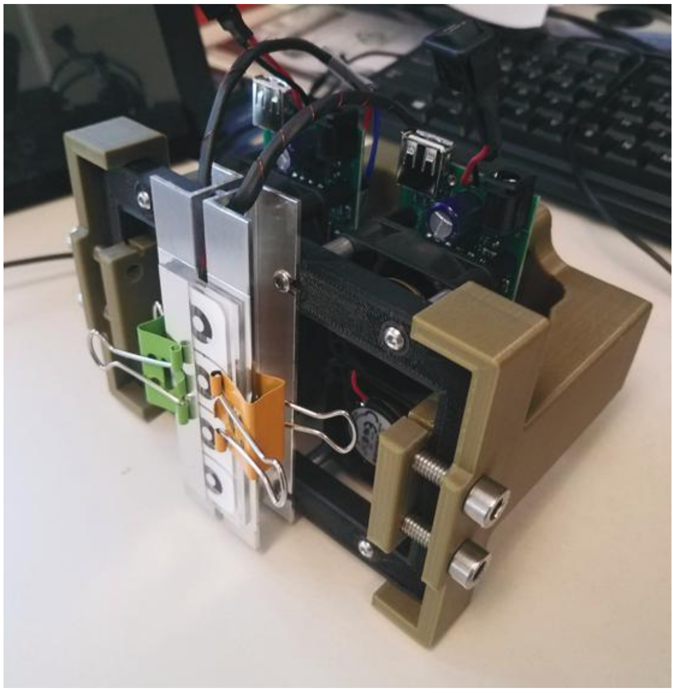

FIG. S6: Thermal unit of the research-grade Donut PCR instrument. **(a)** Schematic of the heating unit with two aluminum heaters and one-chamber chip (reduced in size for better visualization). **(b)** Real-life image of the heating unit with four-chamber chip fixed on it. The heating unit is shown along with heating unit holder.

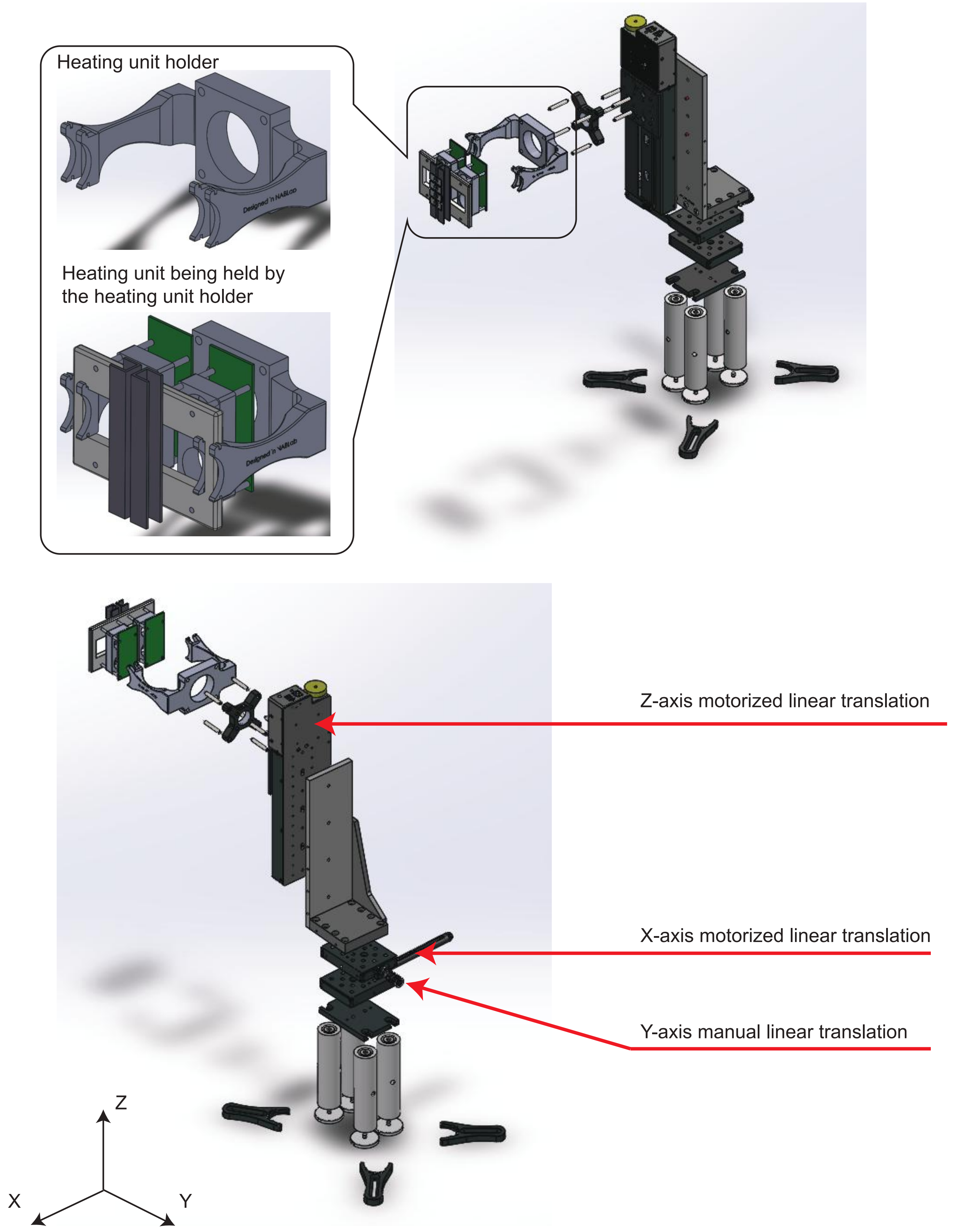


FIG. S7: Schematic of the mechanical system of the research-grade Donut PCR instrument. Shown is "torn apart" instrument comprising motorized Z- and X- electromechanical linear translator stages covering distances of 150 and 25 mm, respectively. Focusing (Y-axis movement) of the chip is realized by a linear stage, that can travel in 25 mm with a resolution of 10 $\mu$m.

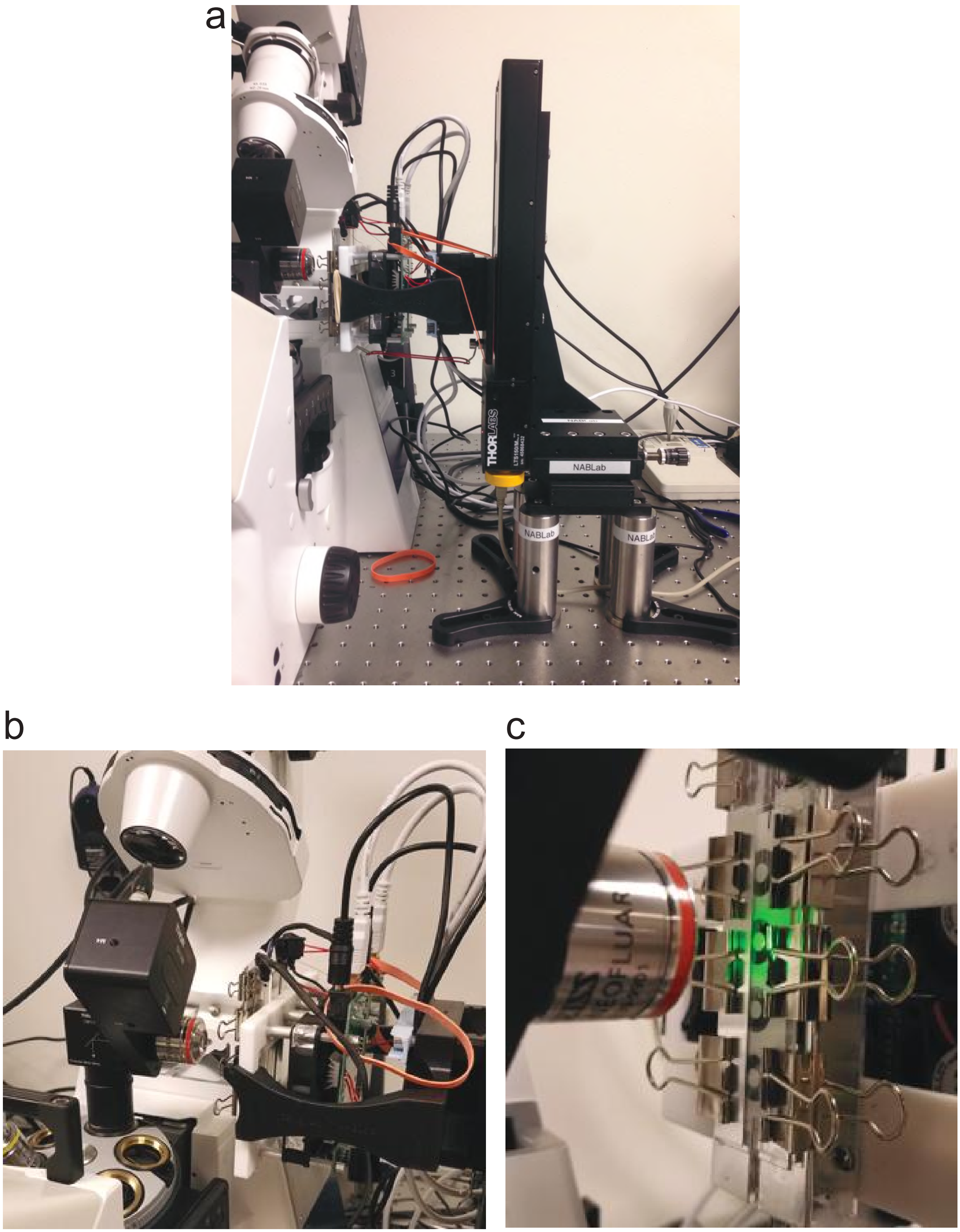


FIG. S8: Real-life images of the research-grade Donut PCR instrument mechanical system. **(a)** Shown here is the entire system. **(b)** Zoomed image of the optical system along with the heater holder. **(c)** Shown here is the 4-chamber Donut PCR chip illuminated with green light via opened optical beam shutter.

### S3. IMAGE PROCESSING AND QUANTITATION

Pipeline for image processing and quantitative data extraction is outlined in Fig. S9. The pipeline consists of three main blocks: raw data collection, image analysis, and data analysis. Collection of fluorescent images was usually performed at 30 or 60 sec intervals. If the analysis was run on a multi-chamber Donut PCR chip, then the collected images were first divided to a number of subfolders corresponding to the number of the fluidic chambers that the Donut PCR cartridge had.

Fig. S10 shows main steps of the script for fluorescent image processing implemented in Matlab. These steps are the same for all assays. After obtaining a set of fluorescent images (Fig. S10a) automatic spot recognition is performed using circular Hough transform (Fig. S10b) [1]. This usually results in correct identification of all microarray spots presented in the image. However, sometimes a few spots could be missed and some spot misidentification could occur, mainly because of low fluorescence intensity of the quenched probes (Fig. S10c). Thus, at the next step of the pipeline, both missed and misidentified spots can be manually added or removed. In addition, during this step, one extra spot, namely - "background" spot (shown in yellow), is also added manually to take into account changes in background fluorescent behavior during the run. The "background" spot has to be positioned in such a way that during the following step of spot sorting and number assignment, the"background" spot is assigned to number #1 (Fig. S10d). Spot sorting is performed in rows from top to bottom, and in columns from left to right. After confirming correct spots sorting, for each identified spot, including the "background", a logical (0,1) mask is generated (Fig. S10e). The step of the logical mask generation represents producing a pseudo image with the same pixel resolution but consisting of only 0s and 1s. Locations of 1s are determined by the area of interest, in our case - the location and the size of a particular spot, while 0s are assigned for the rest of the pixels. Finally, spot intensity is calculated by multiplying each mask by each fluorescent image (digitalized) and averaging the value of all non-zeros pixels (Fig. S10e).

After intensities for all spots are calculated for all images taken, the time-based fluorescence evolution for each spot is calculated (Fig. S11 Step 1). Occasionally, smoothness of the raw data is compromised by ambient light fluctuation, dust appearance on the essential parts of the optical system and inconsistency of the image acquisition hardware/software. In this case, a procedure of raw data processing takes place. Data processing is performed for each spot. It begins with subtracting background fluorescence intensity from the spots intensity to compensate the intensity drift (Fig. S11Step 2.) followed by subtraction of the mean fluorescence values calculated from the first three images (Fig. S11Step 3). If required, intensity normalization to 1 is then performed (Fig. S11Step 4), with the mean spot intensity from the last three images defined to value 1 (max value).

Threshold value can be calculated for each kinetic curve, as shown in Fig. S12a, which can further be used for quantitative analysis of data. As a threshold cut off value, in this study, we used 10% of the entire intensity range for a particular kinetic curve. We showed that using 5% and 15% threshold cutoff values did not significantly change the threshold times ($T_{\mathrm{t}}$) registered (Fig. S12b). However, when calculating $T_{\mathrm{t}}$s at the levels of 20% and 25% of the total fluorescent increase, some outliers appeared. This observation could be explained by that the amplification reaction turned into linear amplification phase after fluorescence signal reached 20% and 25%.

Implementation of the described image processing and data analysis pipelines in Matlab script is provided as supplementary materials and includes exemplary amplification dataset for the whole 20-plex and 4 subsets of primers, as well as a time-lapse video of fluorescence signal accumulation.

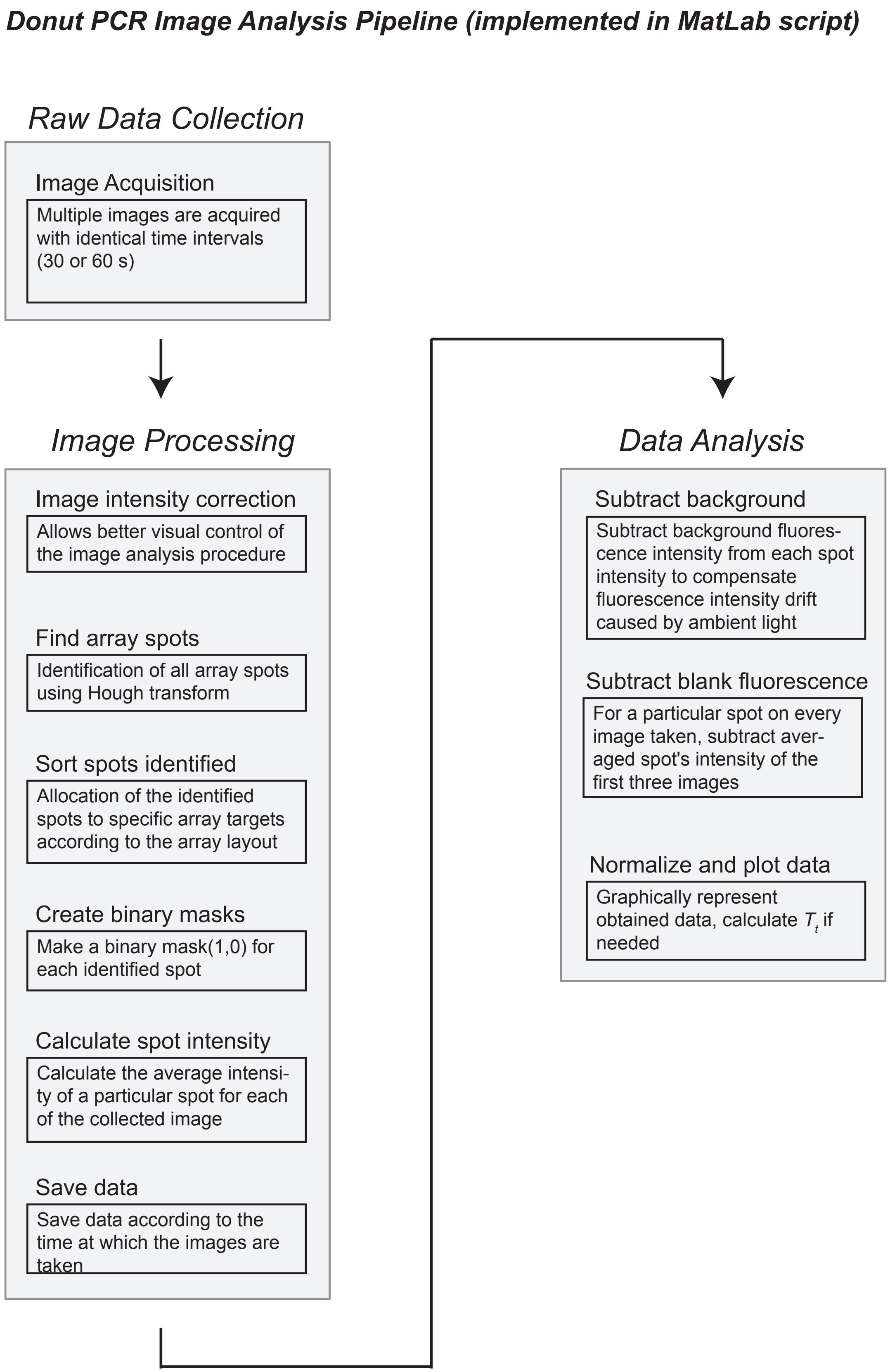


FIG. S9: Data analysis pipeline.

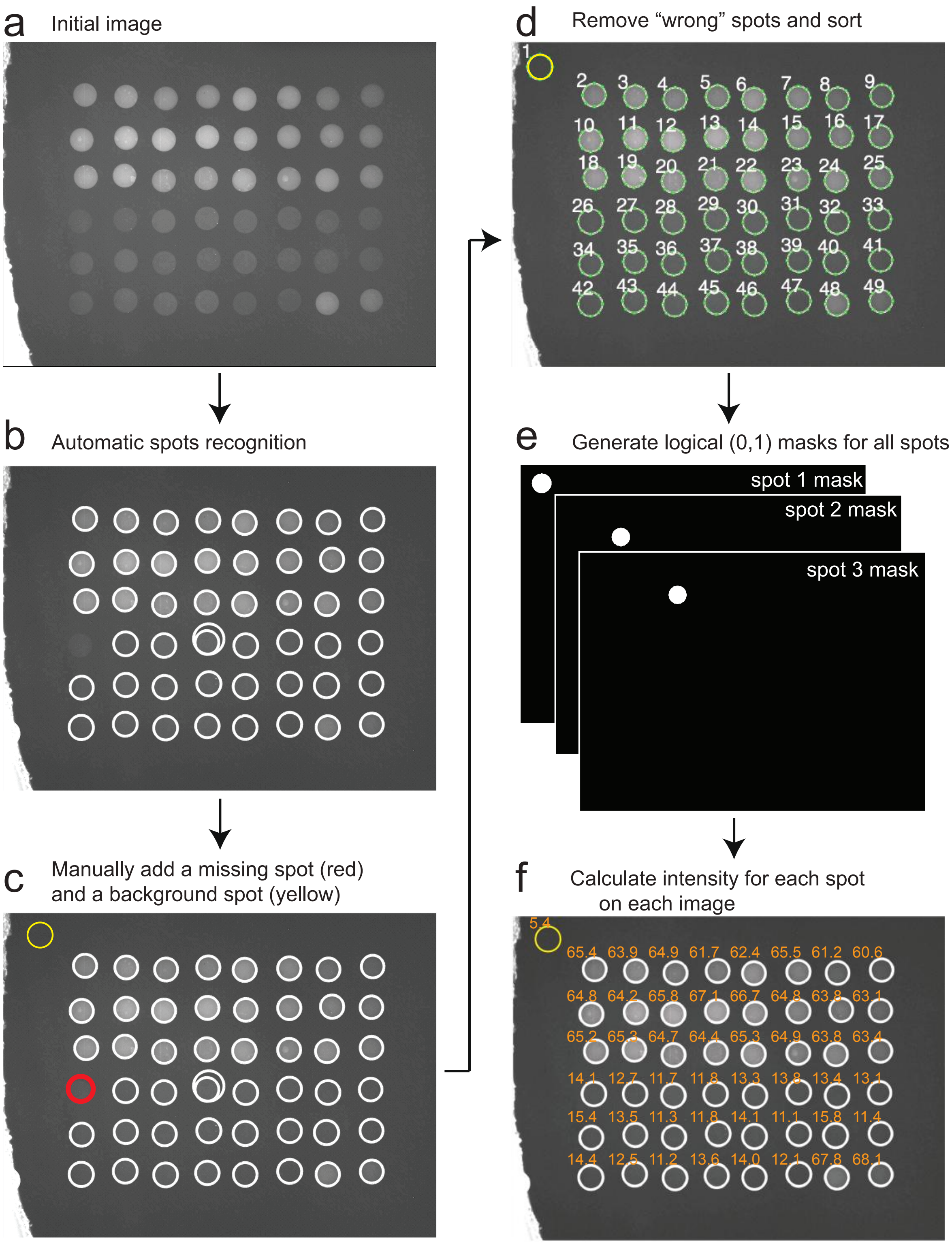


FIG. S10: Fluorescence image processing pipeline.

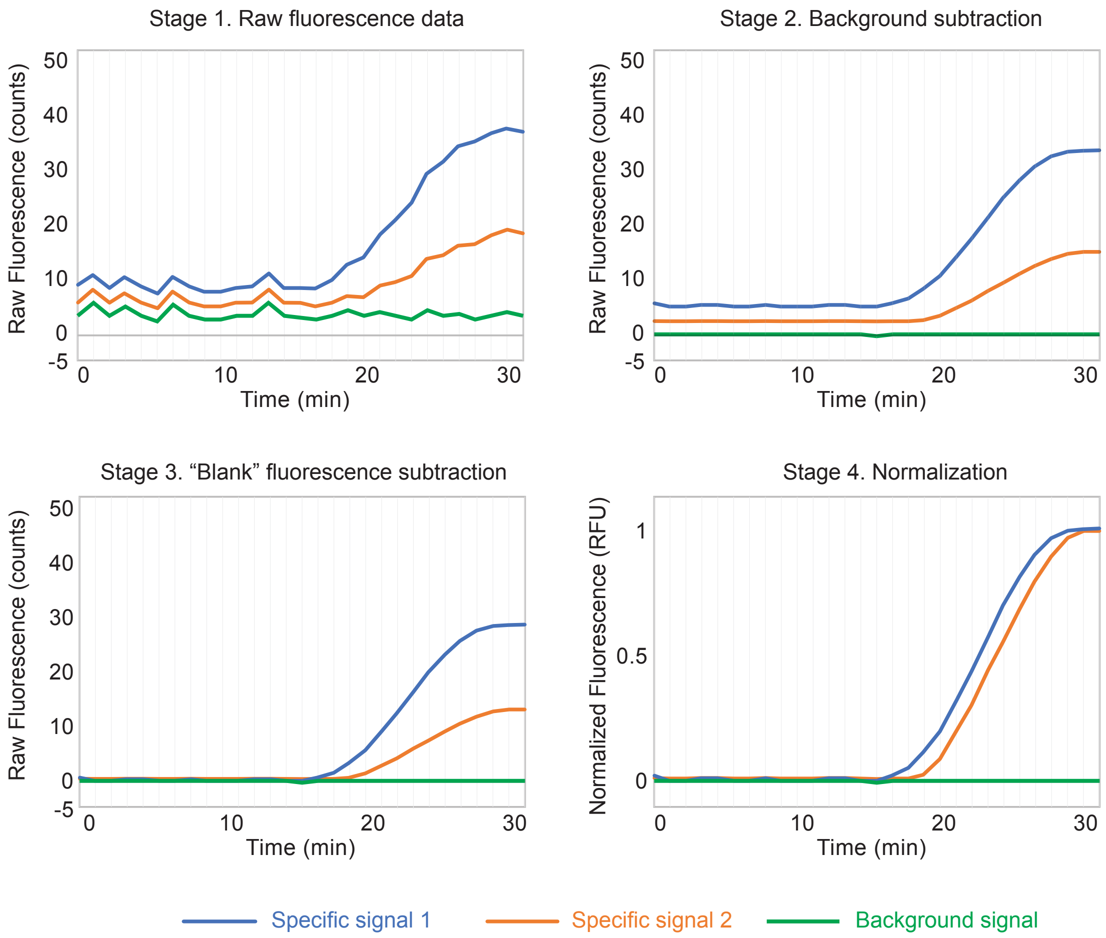


FIG. S11: Pipeline for raw fluorescence data treatment and normalization. **Stage 1** shows raw fluorescence data obtained after fluorescence image processing for two specific (brown and blue traces) and background (green) spots, as an example. Identical intensity spikes and falloffs were observed for all three kinetic curves including background trace. **Stage 2** shows the data after background subtraction. Traces became smooth indicating that intensity spikes and falloffs previously seen had the same origin, for example, from inconsistency in the ambient light of the imaging software. **Stage 3** is a graphical presentation of the data after normalizing to 0 performed by subtraction of the averaged intensity of the first 3-5 cycles, for each spot separately. Finally, **Stage 4** shows scaling the data between 0 and 1, where the average fluorescence intensity of the last three readings was treated as 1.

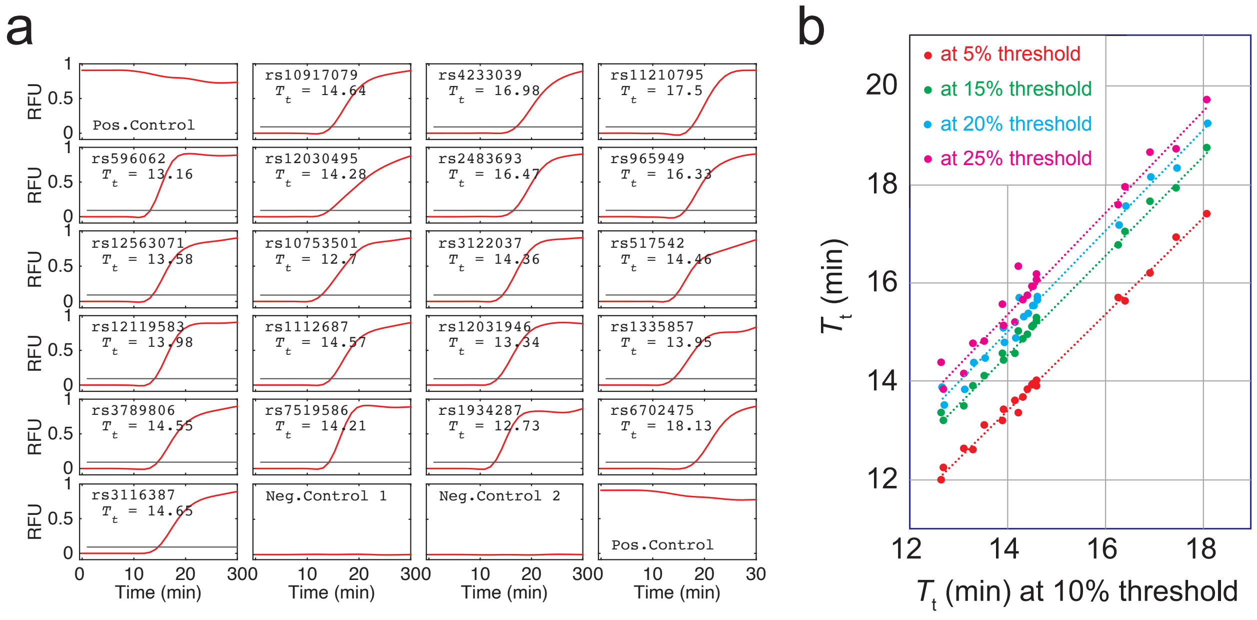


FIG. S12: Selection of the threshold (cutoff) value for threshold time evaluation and quantitative analysis. **(a)** Typical set of 20-plex Donut PCR amplification kinetic curves generated after amplification with 10 ng of human genomic DNA. Threshold time ($T_t$) for each curve was calculated based on a threshold value of 0.1 RFU (10% of the fluorescence intensity range) used as a default in this study. **(b)** Comparison of threshold time distribution when 5%, 15%, 20% , and 25% threshold values were used for $T_t$ calculations.

### S4. POLYMERASE SELECTION

Donut PCR amplification has been successfully shown to work robustly with different types of DNA polymerases. As shown in Fig. S13a, we tested three types of commercially available enzymes: JumpStart Taq from Sigma, Q5 HotStart polymerase from NEB and AptaTaq exo- polymerase from Roche Custom Biotech. All polymerases allowed the generation of specific amplification product without visible primer dimers or nonspecific amplification. However, behavior of the probes showed large differences depending on the DNA polymerase used. Fig. S13b shows kinetic fluorescent trace of the fully quenched probe in the presence of the temperature activated JumpStart polymerase (no primers or dNTPs were added to the reaction). Before adding the polymerase, the probe was stably quenched. After the enzyme was added, the probe's fluorescent intensity started to increase. An explanation for such an observation could be that the 5'→3' exonuclease activity of the enzyme cut the quencher from the probe and caused the fluorescence to increase.

An opposite situation was observed while testing compatibility of the Q5 HotStart DNA polymerase (Fig. S13c). For the unquenched probe, after injection of the polymerase in the reaction chamber, the decay of the fluorescence signal registered from the probe was observed. In contrast to Taq DNA polymerase, Q5 had 3' → 5' exonuclease activity (proofreading activity), that resulted in the detachment of the fluorophore from the probe and as a result - decrease of fluorescence signal. Indeed, the stability of the probe before the Q5 polymerase was added was not compromised.

Based on previously obtained results, we anticipated that a polymerase lacking both 5' → 3' and 3'→5' activity would fit the best Donut PCR assay requirements. As expected, both quenched and unquenched probes, in a solution of AptaTaq exo-DNA polymerase, did not show any significant change in the spot fluorescence intensities (Fig. S13d). Similar results were obtained for Platinum Tfi Exo(-) DNA Polymerase from Invitrogen (data not shown).

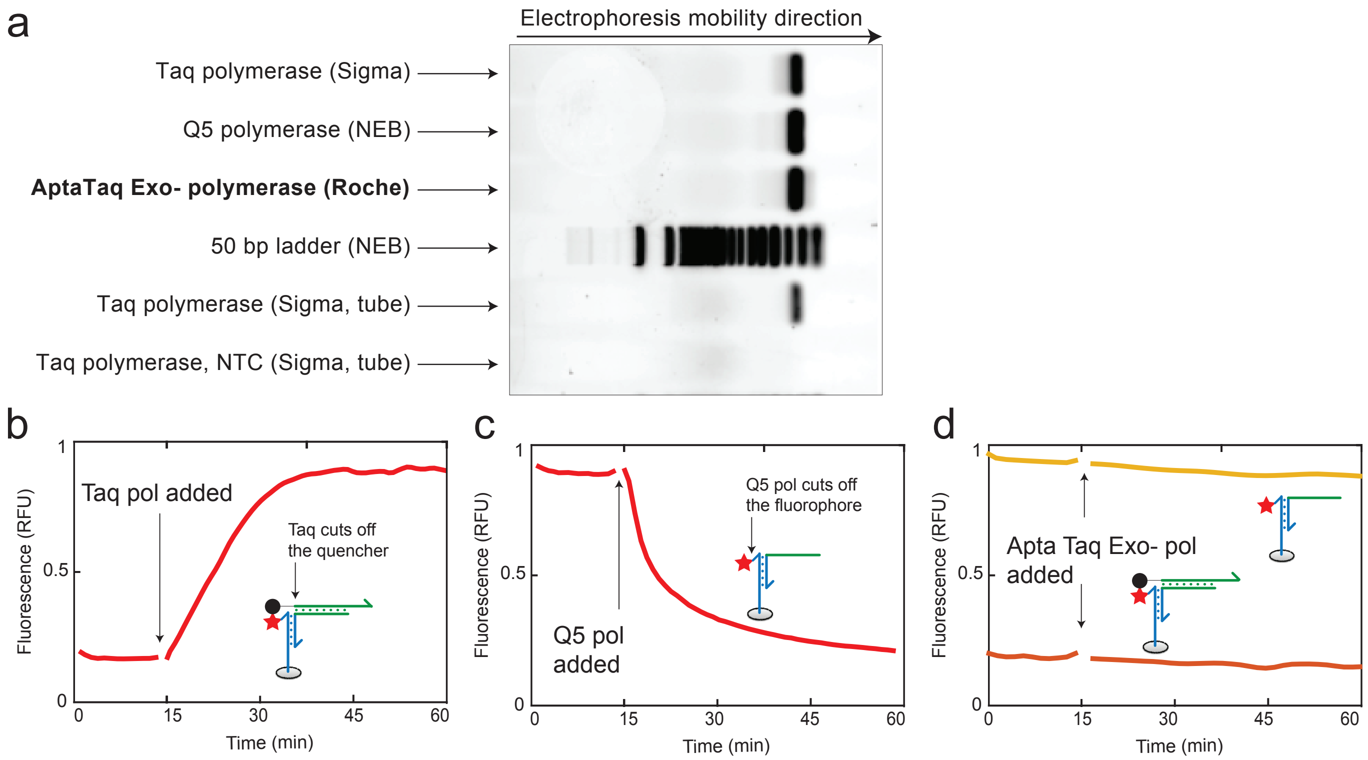


FIG. S13: Selection pf DNA polymerase for robust amplification and detection. **(a)** Electrophoretic evaluation of three DNA polymerases used for Donut PCR amplification. As a control, amplification in a standard Peltier PCR thermocycler was performed. **(b)** Kinetic fluorescence trace of the fully quenched probe (insert) during its incubation in the presence of the Taq polymerase (JumpStart DNA polymerase, Sigma). **(c)** Kinetic fluorescence trace of the un-quenched probe (insert) during its incubation in the presence of the Q5 HotStart DNA polymerase (NEB). **(d)** Kinetic fluorescence traces of both quenched (orange) and un-quenched (yellow) probes in the presence of the AptaTaq exo- DNA polymerase (Roche).

### S5. FLUID DYNAMICS IN DONUT PCR CHAMBER

#### Distribution of flow velocities in the convection flow

The default thickness of the Donut PCR reaction chamber is 360 $\mu$m. This chamber can be considered as a passive microfluidic system with a laminar flow behavior. We measured flow velocities for different chamber thicknesses (see Fig. 2b of the main manuscript). The data were generated from 10 consecutive images with 20 particles per image on average. It was clearly seen that majority of the particles had the speed from 800 to 1200 $\mu$m/s that corresponds to 26 to 17.5 seconds per lap with an average of circulation speed of 22.2 sec/lap. However, a small number of "slow" particles with velocities less that 800 $\mu$m/s (circulation time greater than 26 sec) was observed. This is explained by parabolic velocity profile distribution in microfluidic systems where the flow slows down being closer to surface. The same phenomenon explains a small fraction of "fast" particles (with the speed greater than 1200 $\mu$m/s and circulation time close to 15 sec/lap) circulating in the very middle of the chamber.

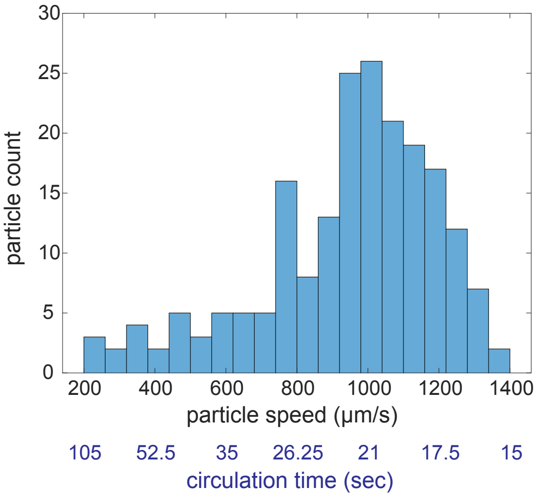


FIG. S14: Velocity distribution of N=200 particles in the 360 $\mu$m chamber heated to 95°C and 60°C. Average circulation time was 22.2±3 sec per lap. Velocities of the particles were measured from 10 images containing roughly 20 particles per image with the exposure time of 0.5 sec.

#### Flow geometry in three different convection flow chambers

Fig. S15 shows expected flow patterns in the three convective flow chambers tested in manuscript Fig. 2a and explains the nonspecific amplification in the "pizza"-shaped chamber and capillary tube. We believe that the presence of 'dead' zones in the middle of the pizza-shaped chamber and capillary tube has minimal convective circulation (yellow tracks). In these zones, primers stay for extended times at low temperatures, facilitating the formation of both primer dimers and nonspecific genomic amplification. In contrast, design of the donut-shaped chamber eliminates all temperature stalled zones and provides correct shuttling of the entire PCR solution between the denaturing and annealing temperature zones.

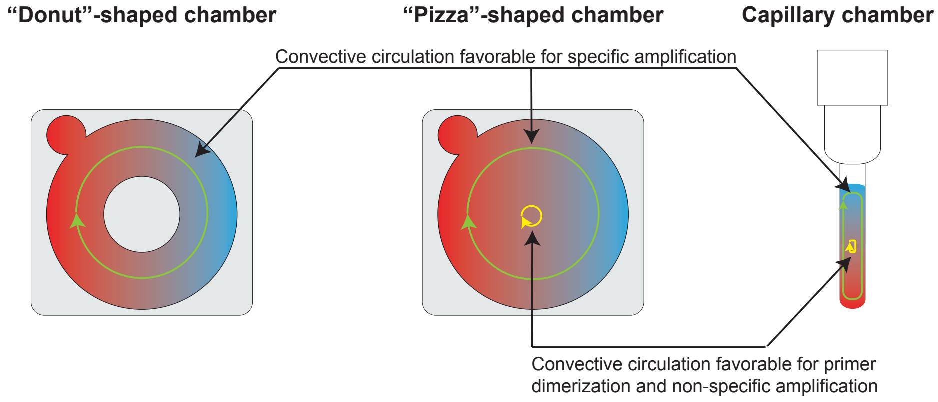


FIG. S15: Schematic representation of the annular flow profiles in three convection flow chambers. The yellow cycles indicate low velocity, low-temperature zones where primer dimer and nonspecific genomic amplification is believed to be frequent.

**Donut PCR amplification speed depends on chamber thickness**

We expected that the circulation speed would have a pronounced effect on the amplification speed and turnaround time. To confirm our hypothesis, we performed amplification of the 200 bp region of the human *TFRC* gene (10 ng input = 2900 haploid genomic copies) in Donut PCR chambers of different thicknesses (Fig.S16). Detection of the amplification product was performed using *TFRC*-amplicon specific probe immobilized on the surface of the reaction chamber. We compared the overall efficacy of amplification/detection by comparing time points when the fluorescence intensity of the probe started rising (Threshold time, $T_t$). For 450 and 360 $\mu$m chambers the $T_t$ were measured to be roughly 13.5 and 14.4 min, respectively. (Fig.S16a). Decreasing the chamber thickness down to 270 $\mu$m resulted in a 7 min delay in $T_t$ compared to the 360 $\mu$m chamber. The thinnest chamber of 180 $\mu$m had no amplification detected within the first 30 min. Fig. S16b shows nearly perfect correlation ($R^2$ = 0.99) between the observed speed velocity, measured by particles tracking, and amplification $T_t$ values.

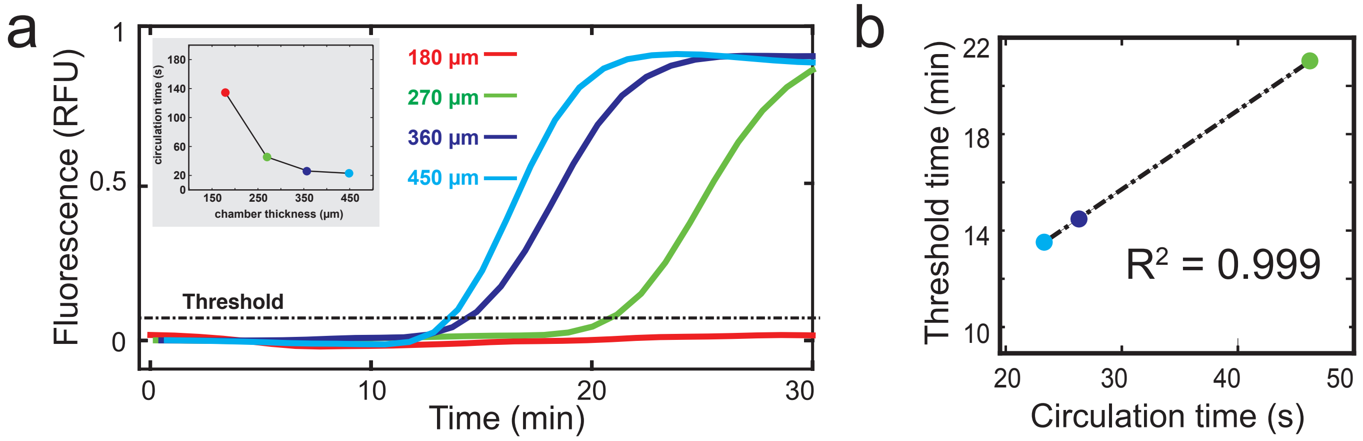


FIG. S16: Dependence of the amplification speed on the Donut PCR chamber thickness, based on convection flow velocity. **(a)** Kinetics of fluorescence signal accumulation while amplifying and detecting human *TFRC* gene in the Donut PCR chamber of different thicknesses. **(b)** Correlation between convection flow circulation time and threshold time $T_t$ for chamber thicknesses of 450 (light blue), 360 (blue) and 270 $\mu$m (green).

### S6. QUANTITATION OF DNA TARGETS BASED ON $T_T$

Fig. S17 shows the amplification curves summarized in the Fig. 2e of the main manuscript. In all experiments, the input DNA contained a mixture of human gDNA and mouse gDNA. For better visual comparison, kinetic traces of fluorescence signal accumulation were split according to the specific targets for detection of mouse-specific targets (*Nono* gene), and for detection of human-specific targets (*TFRC* gene). All aliquots of nucleic acids were prepared in 10 mM Tris-HCl solution (pH 7.5) with 0.01% Tween-20 added to mitigate DNA loss from plasticware adsorption.

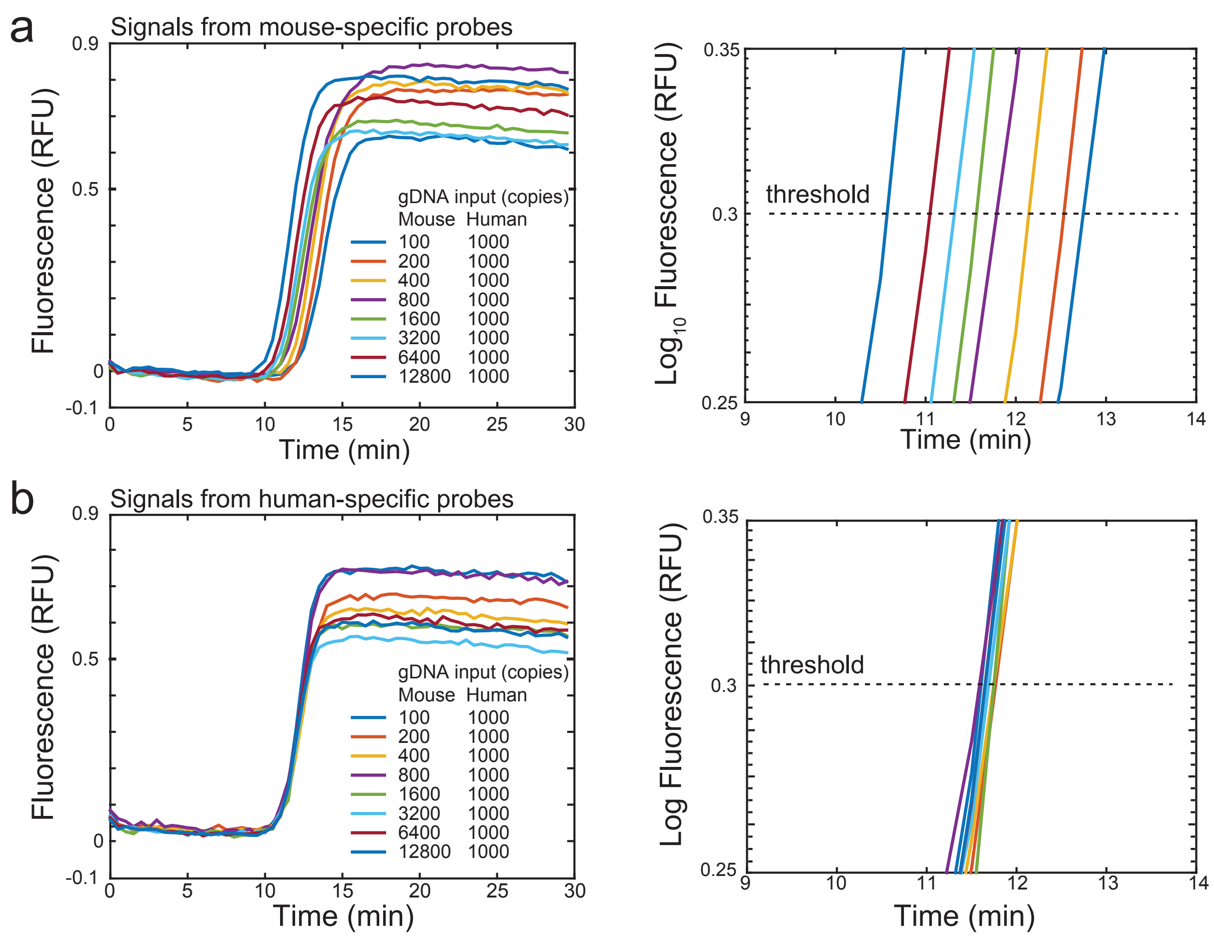


FIG. S17: Amplification curves for nucleic acid quantitation experiments summarized on Fig. 2e of the main manuscript. **(a)** Donut PCR kinetic curves for the mouse *Nono* gene probe. **(b)** Donut PCR kinetic curves for the human *TFRC* gene probe.

### S7. SIMULTANEOUS AMPLIFICATION AND DETECTION OF 20 GENETIC LOCI FROM HUMAN GENOMIC DNA

To demonstrate the highly multiplex DNA analysis, we designed an assay for simultaneous amplification and detection of 20 genetic loci from human genomic DNA. For this assay, 20 independent pairs of primers are designed and used with a chip printed with 20 different target-specific probes. Fig. S18a shows the layout of the array. Each spot on the array is duplicated to assess the reproducibility of the detection. Besides specific spots, the array comprises both positive (PC) and negative (NC) control spots used for grid alignment during image processing and assessment of nonspecific real-time detection, respectively. Amplification was performed along with with all 20 primers using 4 ng (1160 copies) of human genomic DNA as an amplification input. To better show the robustness of Donut PCR amplification and detection we additionally split 20-plex primer set into four subsets in the following way: subset 1 - primers specific to targets 1-11, subset 2 - primers specific to odd numbered targets, subset 3 - primers specific to even numbered primers, and subset 4 - primers specific to targets 12-20. Fig. S18b presents summary of $T_{\mathrm{t}}$ values obtained after amplification with all primers and four primer subsets. Fig. S18c through S19g shows end-point ($30^{th}$ minute of the reaction) fluorescence images of the corresponding assay along with traces tracking accumulation of fluorescence signal from each pair of specific spots (traces are shown as an average of two kinetic traces). It is clearly seen that all spots, where the corresponding primers were not included in the amplification mixture, remained dark, confirming high specificity of the surface probe detection. For targets 5, 7, 12, 16 and 20, measured $T_{\mathrm{t}}$ values remained in 30 s range interval, confirming robust amplification and detection regardless of total number of targets being amplified.

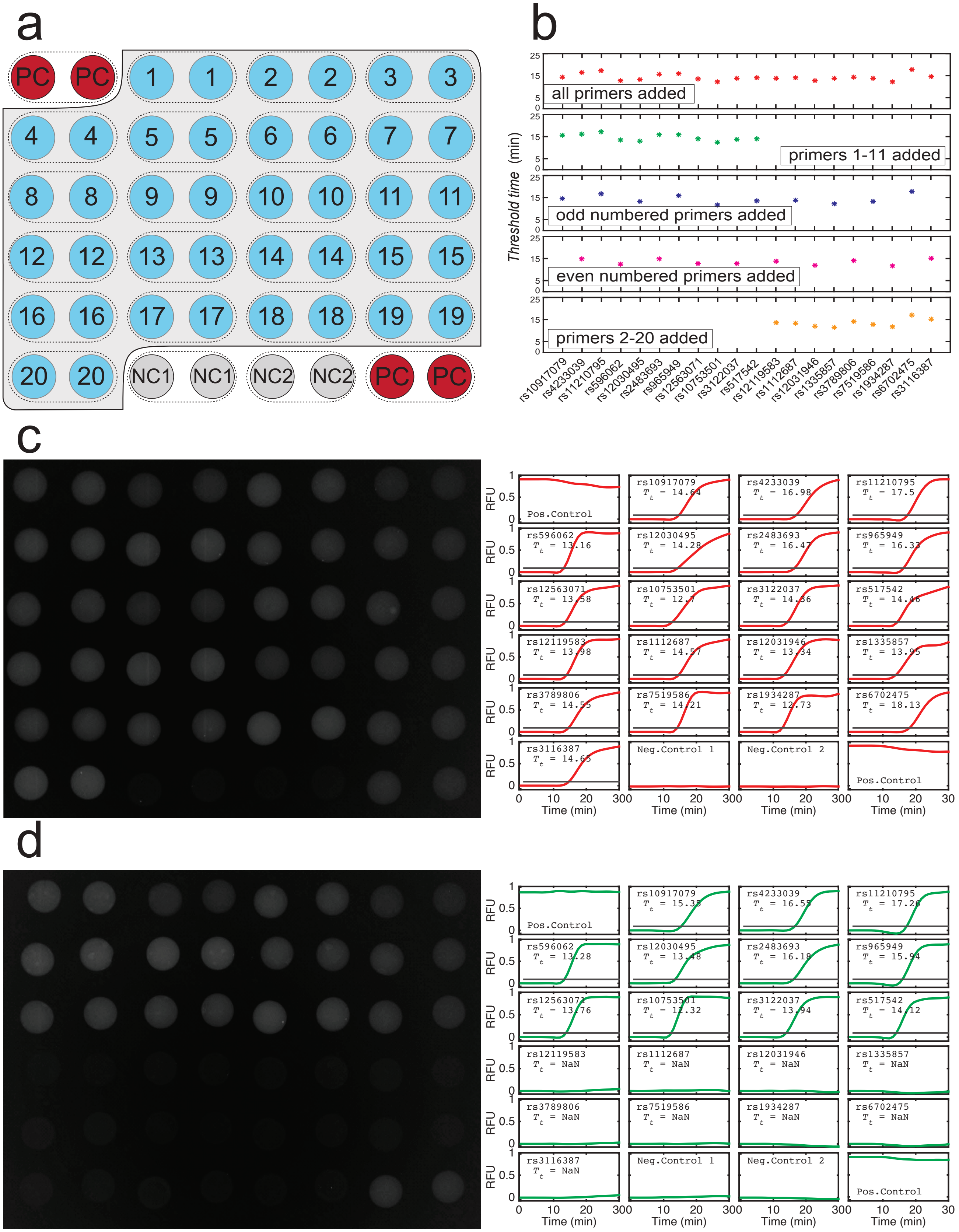


FIG. S18: Simultaneous amplification and detection of 20 human DNA targets using Donut PCR, using 4 ng input gDNA. **(a)** Layout of the Donut PCR probe array. **(b)** Summary of threshold times $T_{\mathrm{t}}$ for experiments where different subsets of primers were added to the reaction mixture. **(c)** End-point fluorescence image after amplification with a full set of 20 primer pairs. **(d)** End-point fluorescence image after amplification with a subset of primers specific to targets 1 - 11.

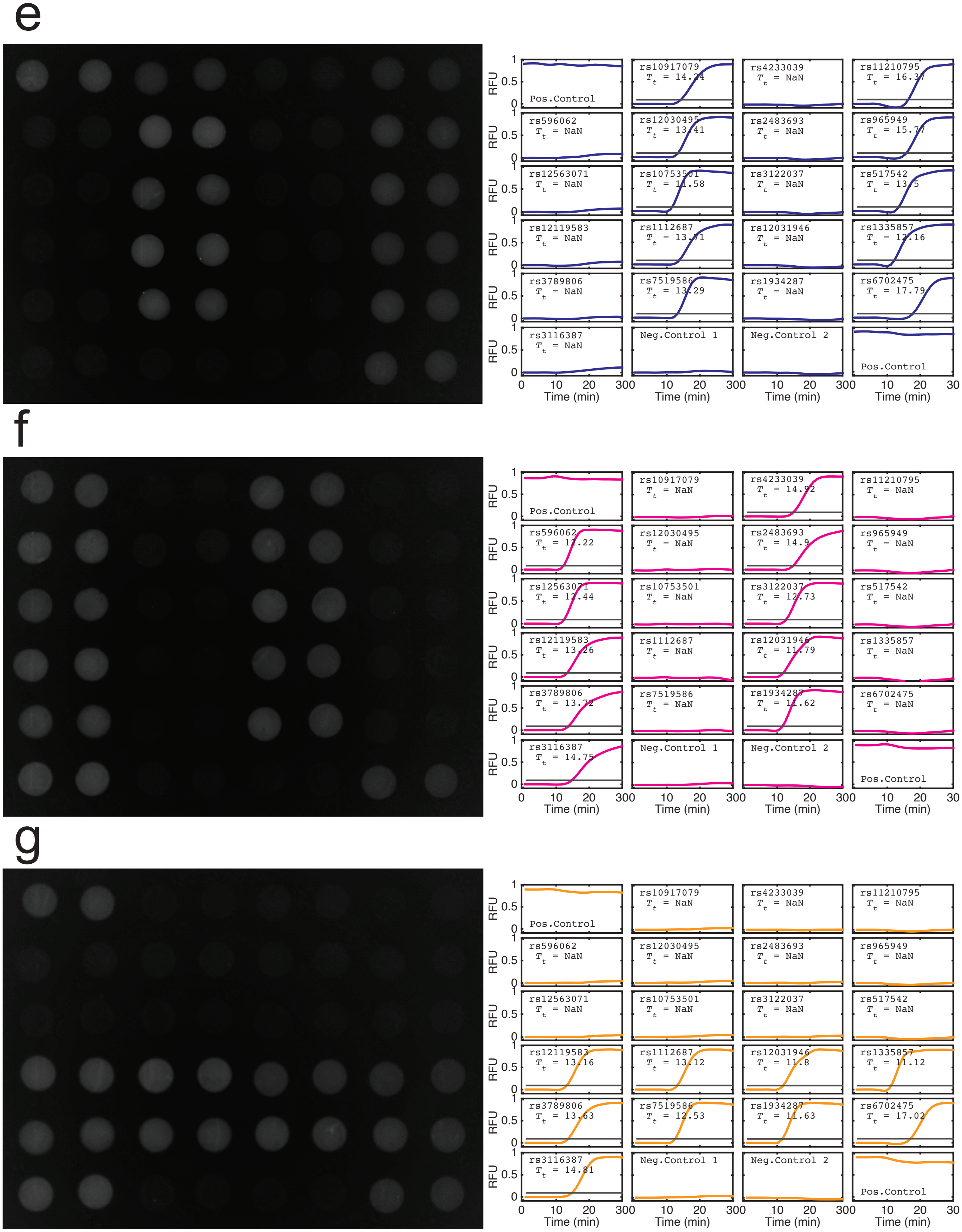


FIG. S19: Continuation of Fig. S18. **(e)** End-point fluorescent image after amplification with a subset of primers specific to odd numbered targets. **(f)** End-point fluorescent image afer amplification with a subset of primers specific to even numbered targets. **(g)** End-point fluorescent image after amplification with a subset of primers specific to targets 12-20.

### S8. BATTERY-POWERED BREADBOARD DONUT PCR INSTRUMENT

The battery-powered breadboard Donut PCR instrument was made using off-the-shelf available components in conjunction with custom-made aluminum heaters and 3D printed parts.A list of the main essential components, including source of purchasing and catalog number is provided as the supplementary Excel file: Components-List-Battery-Powerd-Device.xlsx. Temperature control of the heaters is performed using proportional-integral (PI) algorithm encoded into Arduino Mega 2560 circuit board. Four T-type thermocouples are used as temperature sensors for four heaters (one for each heater) (Fig. S21a). The sensors are embedded into the aluminum plate directly underneath the place where the Donut fluidic chamber is located, and connected to the Arduino board via signal amplifiers. The frequency of data point reading and temperature adjustment is chosen to be 2 Hz. Fig. S21b shows the rate of reaching desired temperature for the heaters powered by a 24 V battery. It requires 5 min to preheat the system with a very short time for temperature stabilization using the chosen temperature control algorithm. One of the heaters has a machined opening to allow image acquisition S20.

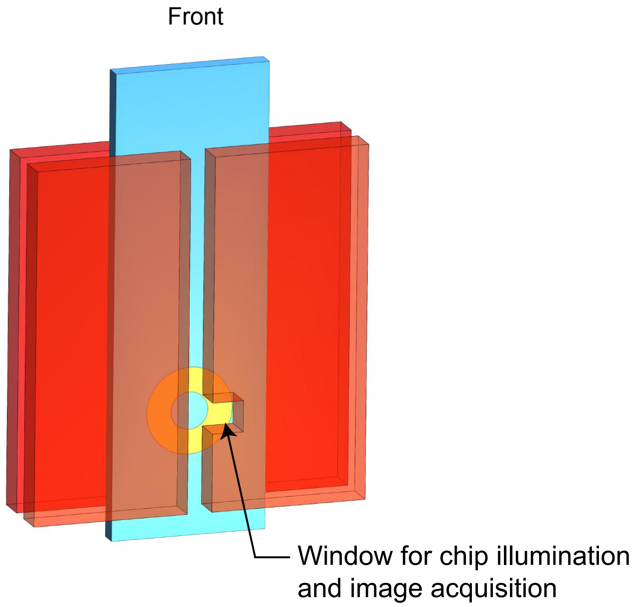


FIG. S20: Schematic of heaters (red) with Donut PCR chip (cyan). The heaters on the left are heated to 95 °C during operation, and the heaters on the right are heated to 60 ° during operation. One of the 60°C heaters has a machined window that allows for optical imaging.

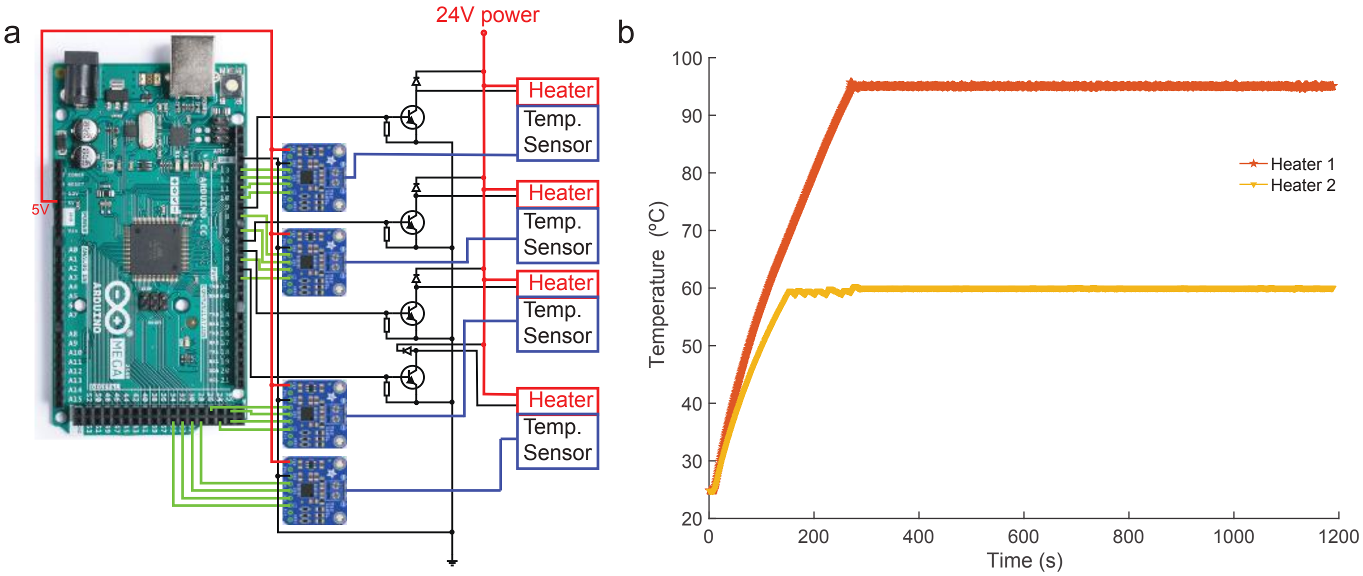


FIG. S21: Temperature controller for the battery-powered Donut PCR device. **(a)** Electrical diagram for the temperature controller implemented on the Arduino Mega 2560 circuit board. **(b)** Kinetic traces of the heaters' temperatures targeting temperature-setpoints of 95 °C and 60 °C.

### S9. BACTERIAL IDENTIFICATION PANEL

#### Design consideration for 16S-based bacterial identification

Our multiplex bacterial identification assay is based on amplification of the 16S rRNA gene using a single pair of conserved PCR primers. The different amplicons insert sequences generated from different bacterial species are detected by different surface-bound probes. To design the primers and probes, we first aligned the 16S rRNA gene from different bacterial species using Genious Prime software and the MUSCLE alignment algorithm (Fig. S23). Hyper-variable region V3, spanning nucleotide positions 400-600 of the bacterial 16S rRNA gene, was chosen as the target sequence region. The universal 16S rRNA primers for this assay generates a ≈200 nt amplicon.

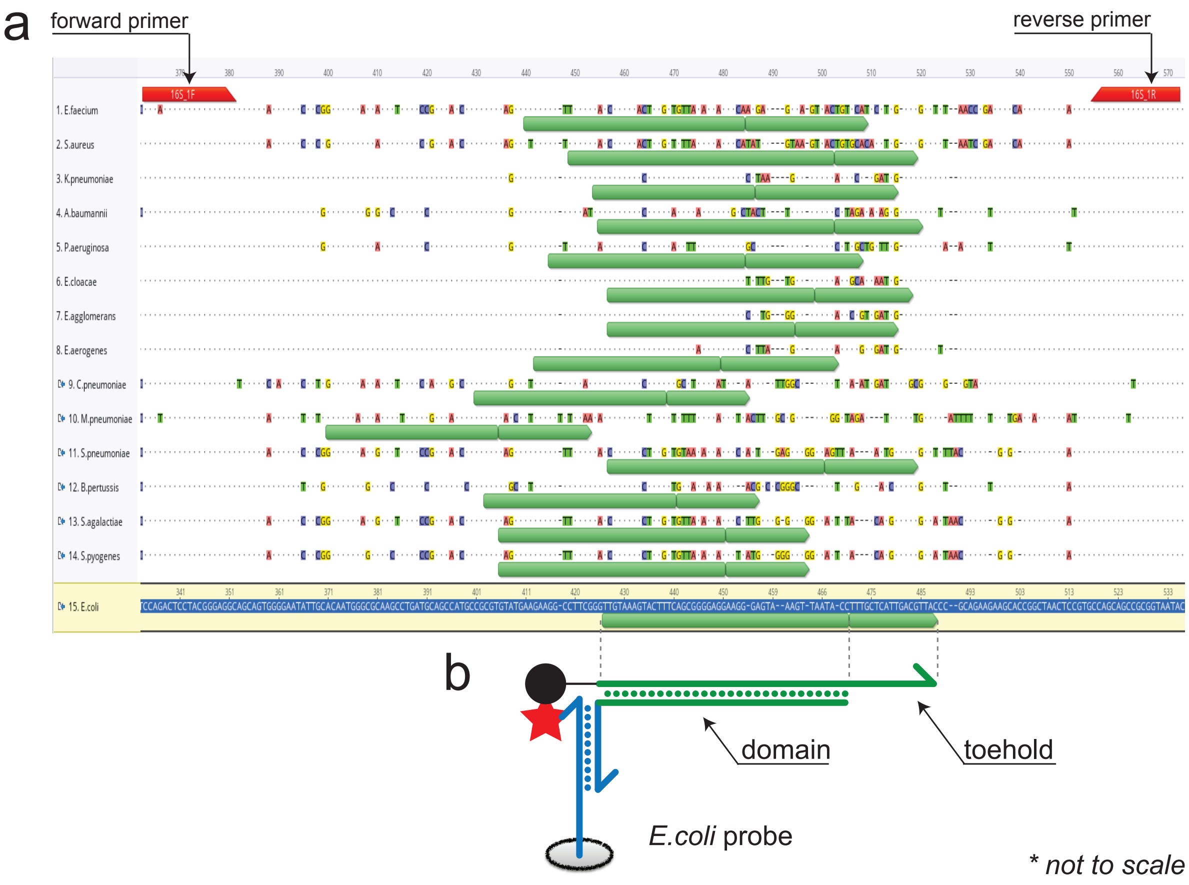


FIG. S22: Schematic of the 15-plex bacterial identification assay's primer and probe alignment. **(a)** The alignment uses *E.coli* 16S rRNA gene sequence as reference. All dots depicted in alignment indicate the same nucleotide at this particular position as the one in the reference sequence (*E.coli*), while highlighted capitalized letters show discrepancy between the sequences. Locations of the primers are shown in red. Sequences to which the probes are designed to be complementary to are highlighted in green and are split in two parts - toehold and domain (for easier probe visualization). **(b)** Graphical representation (not to scale) of the toehold probe architecture and how it aligns to the *E.coli* amplicon sequence.

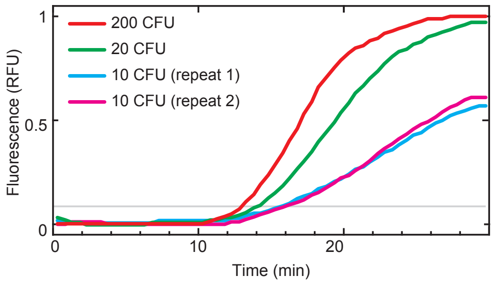


FIG. S23: Donut PCR amplification and detection of low quantities of the *E.cloacae* bacterial gDNA. Serial dilutions of bacterial genomic DNA were made in 1×TE buffer supplied with 0.01% of Tween-20. We were able to successfully detect and discriminate down to 10 colony forming units (CFU), equivalent to 70 copies of the rRNA gene target.

### Detection of 6 bacterial species' DNA using battery-powered Donut PCR instrument

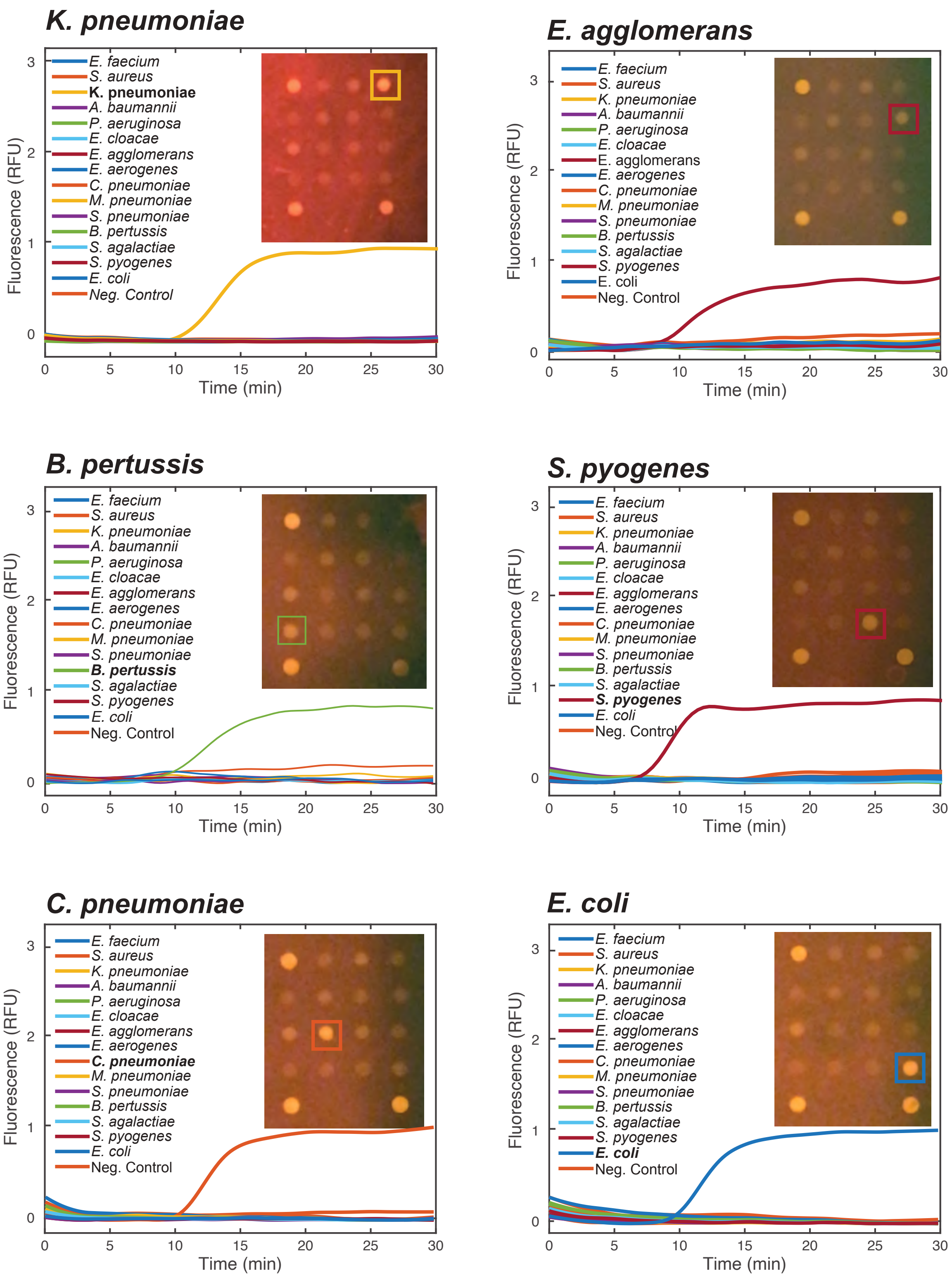


FIG. S24: Detection of 6 different pathogenic bacteria's DNA using the breadboard battery-powered Donut PCR instrument. 0.5 to 1 ng of purified genomic DNA was used as DNA input. Note that the input sample for *C. pneumoniae* was a mixture of the *C. pneumoniae* DNA and DNA from HEp-2 cells (as sold by ATCC). The intracellular pathogen *C. pneumoniae* is usually cultivated in host cells (HEp-2), thus extraction of the *C. pneumoniae* DNA was accompanied by co-extraction of host DNA. Thus, the *C. pneumoniae* input sample also contained human gDNA.

### Detection of 15 bacterial species' DNA using functional prototype Donut PCR instrument

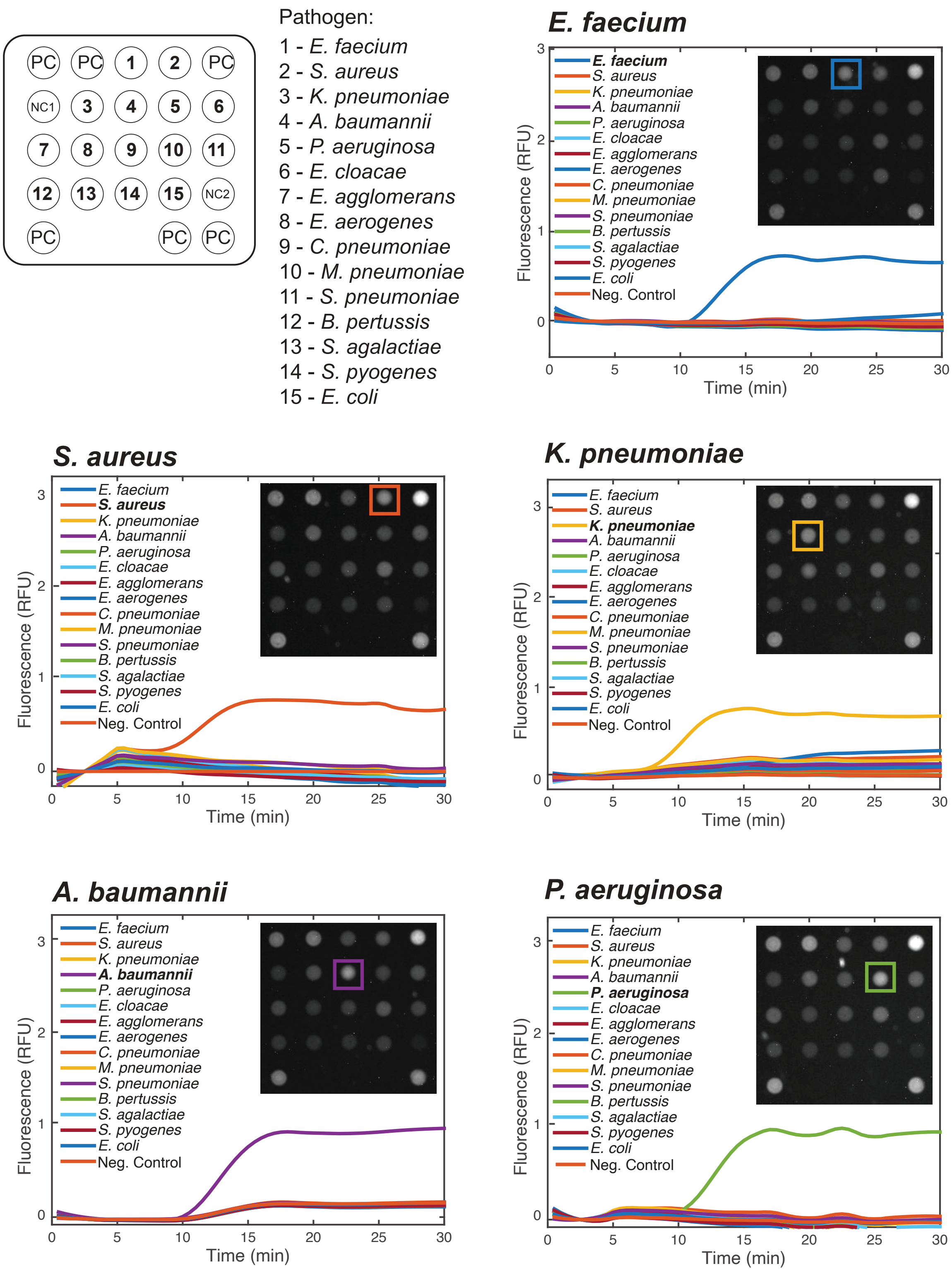


FIG. S25: Detection of 15 different pathogenic bacterial DNA using functional Donut PCR instrument prototype. In the top left corner, we present the layout of the probe array. The remaining subfigures show the fluorescence kinetics and endpoint fluorescence images of Donut PCR assays with different input bacterial DNA. In all cases, threshold time for amplification of 0.5 ng purified genomic DNA was below 12 min. No false positives were detected from nonspecific probe binding of other species' amplicons.

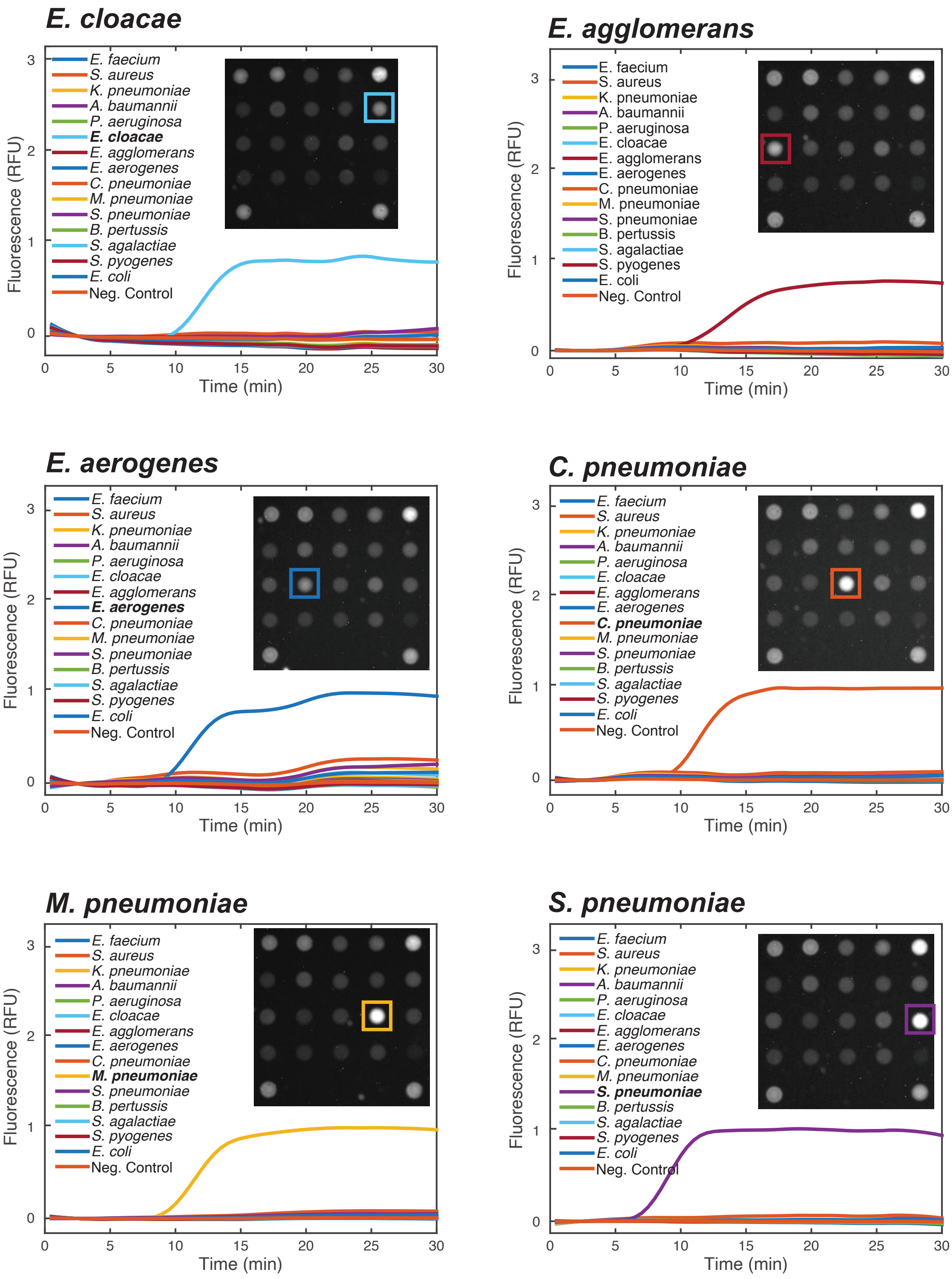


FIG. S26: Continuation of Fig. S25. Input sample *C. pneumoniae* contains also human DNA (see caption of Fig. S24).

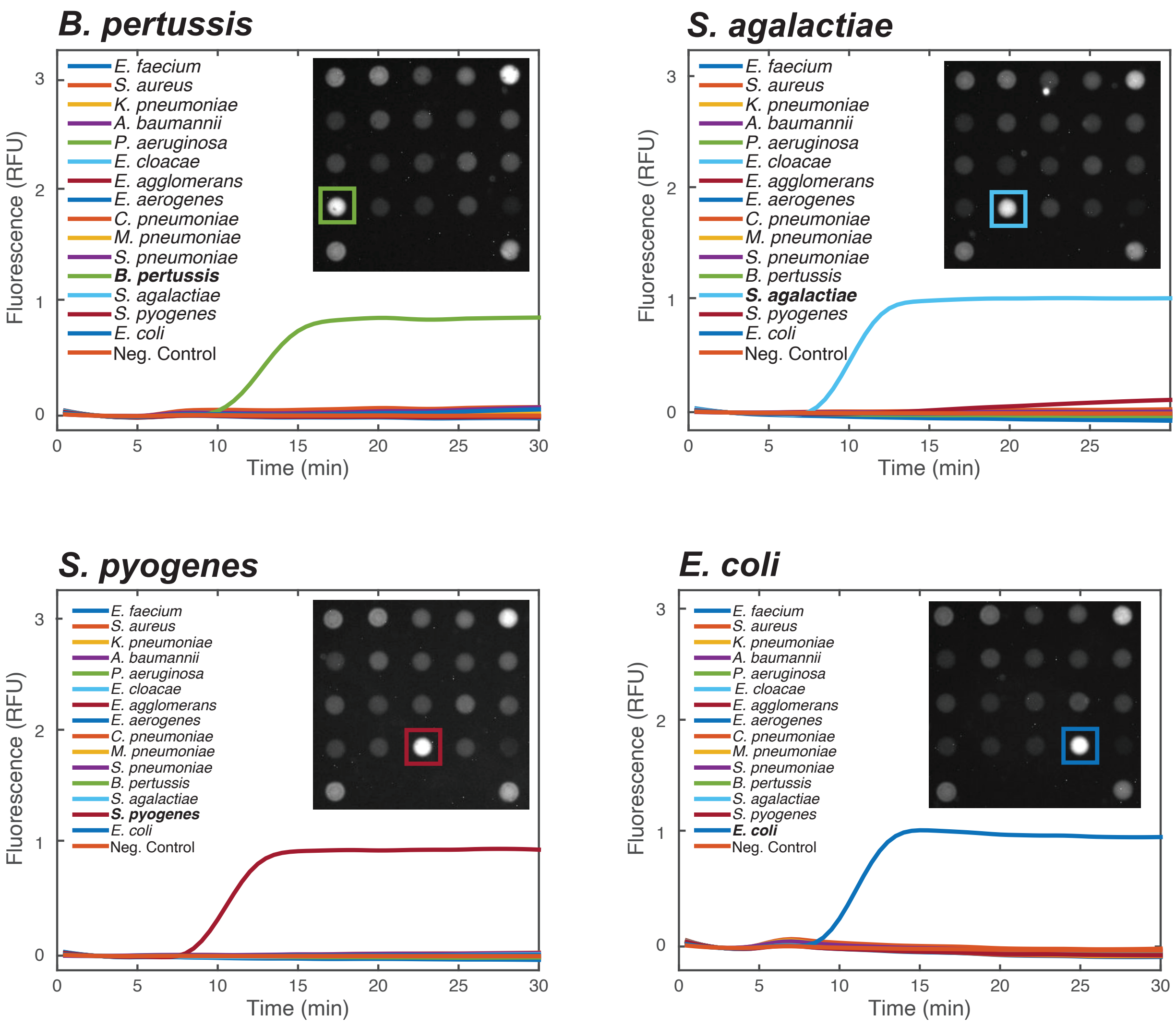


FIG. S27: Continuation of Fig. S25 and S26.

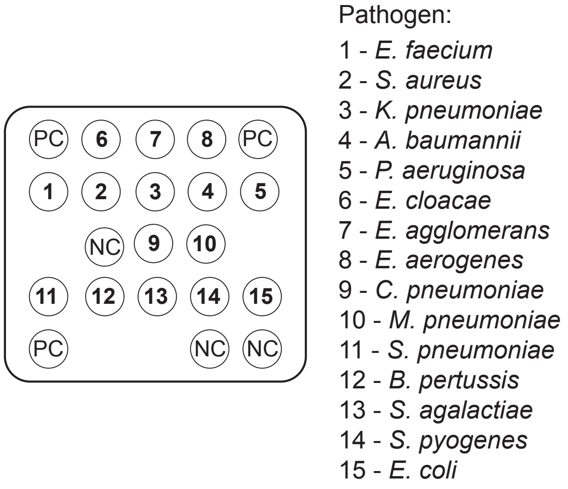


FIG. S28: Layout of the bacterial identification panel presented in manuscript Fig. 4f.

### S10. SINGLE NUCLEOTIDE POLYMORPHISM (SNP) GENOTYPING.

Pairs of toehold probes [2] were designed to detect alternate alleles at each SNP locus. For all probes, the standard free energy of toehold binding was designed to be in a range of -9.5 to -10.5 kcal mol$^{-1}$. For all probes, non-homologous regions (colored in magenta) were designed with standard free energy of binding of approximately -5 kcal mol$^{-1}$. The non-homologous region provides higher specificity by allowing thermodynamic-based discrimination in addition to kinetics-based discrimination of SNP alleles. The phenotypes correlated to the SNP genotypes are listed in Fig. S29a, and Fig. S29b shows the probe layout on the Donut PCR chip.

We first validated the panel on three human reference DNA samples from with well-documented genotypes on the 1000 Genomes project (Fig. S30a). Genotype calls were made based on the threshold time $T_{\mathrm{T}}$ in the fluorescence kinetics curves (Fig. S30b). For this panel, the threshold time $T_{\mathrm{T}}$ was determined based on the time to reach 0.5 RFU, because some SNP loci (e.g. rs17145738) had an initial linearly increasing fluorescence, which may be due to poor probe oligo synthesis. A SNP locus was called as homozygous if the difference in threshold times exceeded 5 minutes for the probes to the alternate alleles. Using this algorithm, all SNP loci were correctly genotyped for all 3 samples.

a

| Target # | Test SNPs | Phenotypes | Mapped Gene | Chr | Alleles |
|---|---|---|---|---|---|
| 1 | rs1558902 | Body mass index | *FTO* | 16 | T/A |
| 2 | rs9939609 | Body mass index | *FTO* | 16 | T/A |
| 3 | rs6265 | Body mass index | *BDNF* | 11 | C/T |
| 4 | rs10889353 | Triglyceride | *ANGPTL3* | 1 | A/C |
| 5 | rs17145738 | Triglyceride | *MLXIPL* | 7 | C/T |
| 6 | rs2954029 | Triglyceride | *TRIB1* | 8 | A/T |
| 7 | rs646776 | Cholesterol | *SORT1* | 1 | C/T |
| 8 | rs12654264 | Cholesterol | *HMGCR* | 5 | A/T |
| 9 | rs4939883 | Cholesterol | *LIPG* | 18 | T/C |
| 10 | rs1532624 | Cholesterol | *CETP* | 16 | C/A |
| 11 | rs10811661 | Blood glucose | *CDKN2A/B* | 9 | T/C |
| 12 | rs1799884 | Blood glucose | *GCK* | 7 | C/T |
| 13 | rs2191349 | Blood glucose | *DGKB-TMEM195* | 7 | G/T |
| 14 | rs1173771 | Blood pressure | *NPR3* | 5 | G/A |
| 15 | rs3827760 | Hair thickness | *EDAR* | 2 | A/G |

b

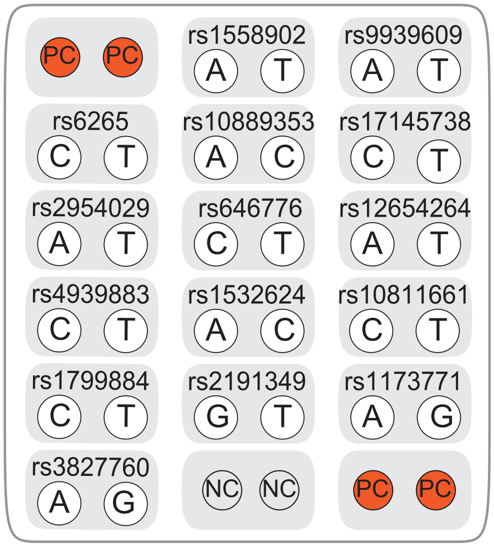


FIG. S29: **(a)** List of human SNP loci included in the SNP genotyping panel. The panel includes a total of 30 probes targeting the alternate alleles of 15 SNP loci. **(b)** Layout of the Donut PCR probe panel.

a

| Target # | Test SNPs | Genotypes of reference DNA samples from 1000 genome project NA18596 | NA18537 | NA18625 |
|---|---|---|---|---|
| 1 | rs1558902 | T/T | T/T | T/T |
| 2 | rs9939609 | T/T | T/T | T/T |
| 3 | rs6265 | T/T | T/T | C/T |
| 4 | rs10889353 | A/A | A/A | A/C |
| 5 | rs17145738 | C/C | C/T | C/C |
| 6 | rs2954029 | A/T | T/T | T/T |
| 7 | rs646776 | T/T | T/T | T/T |
| 8 | rs12654264 | A/T | A/T | A/A |
| 9 | rs4939883 | C/C | C/T | C/C |
| 10 | rs1532624 | A/C | C/C | C/C |
| 11 | rs10811661 | C/T | T/T | C/T |
| 12 | rs1799884 | C/T | C/T | C/C |
| 13 | rs2191349 | T/T | G/T | T/T |
| 14 | rs1173771 | A/G | A/G | A/G |
| 15 | rs3827760 | G/G | G/G | G/G |

b

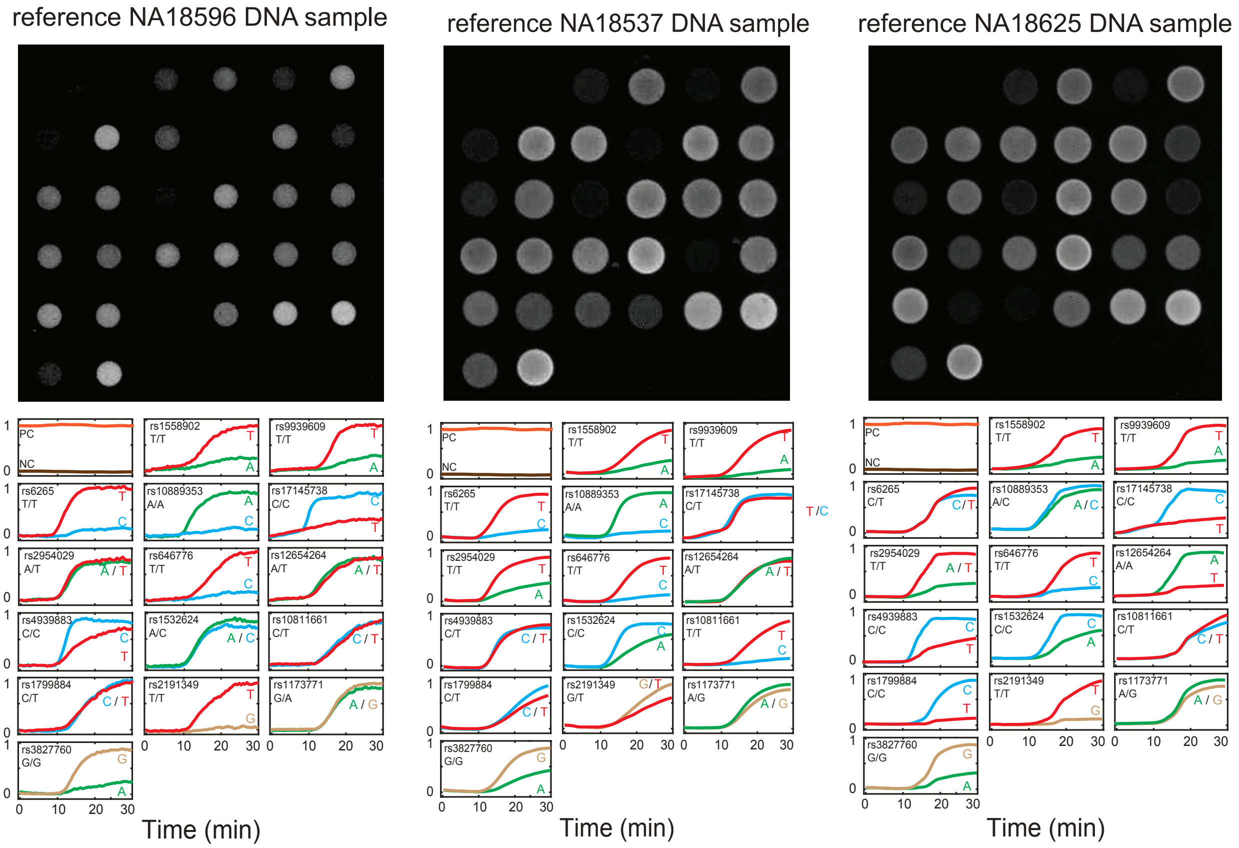


FIG. S30: **(a)** Reported genotypes (1000 Genomes) of the NA18596, NA18537, and NA18625 human gDNA samples (Coriell) used for panel validation studies. **(b)** Endpoint and fluorescence kinetics results of the panel on the NA18596, NA18537, and NA18625 human gDNA samples (8 ng input). The endpoint images shown here are obtained by subtracting the first images (t = 0 min) from the final images (t = 30 min), so the positive control spots do not show up. Fluorescence kinetics for the positive and negative control spots are presented on the same plot for comparison.

#### SNP genotyping using buccal swabs

We next explored the use of whole biological samples with minimal sample processing as input for the Donut PCR system. Here, the DNA samples are released from buccal swab cells using thermal and chemical treatment, with proteins denatured using proteinase K. We used QuickExtract DNA Extraction Solution (Lucigen) to treat buccal cells collected with sterile buccal swabs (Isohelix). The swab was immersed in the QuickExtract solution and heated at 60 °C for 10-15 min to release DNA from cells and then at 95 °C to inactivate the extraction reagents. After the second heating step, the lysate was ready to be used as a DNA input for SNP genotyping. Fig. 5d from the manuscript on genotyping a family trio uses this sample type and approach. The fluorescence kinetics traces are shown in Fig. S31.

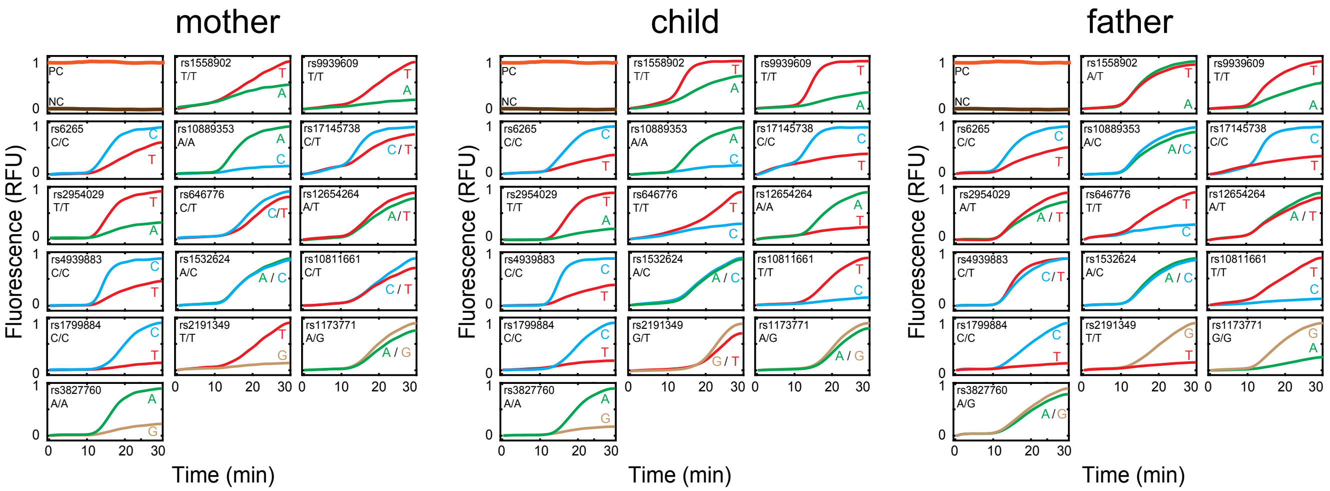


FIG. S31: Time-based fluorescent plots of the SNP genotyping assay performed for three members of one family. DNA input was 2$\mu$L of buccal cell lysate obtained using QuickExtract DNA Extraction Solution (Lucigen). No additionally steps of DNA extraction/purification and quantitation of nucleic acids were performed prior to the genotyping.

### S11. DONUT PCR CAPABILITIES FOR FURTHER DEVELOPMENT.

#### Ultrafast Donut PCR

Given the observation that thicker chambers allow faster convection-based fluid circulation, which in turn leads to faster assay turnaround time, we next designed a 700 $\mu$m-thick chamber to further improve the Donut PCR turnaround time for point-of-care applications. For these experiments, we used the SYTO-82 intercalating dye to report DNA amplicon buildup, to separate the probe hybridization time from the PCR amplification time. To prevent the primer binding from being the kinetic bottleneck to PCR amplification, we also increased the concentrations of the primers from 200 nM to 2 $\mu$M. se reagents. Because the concentrations of the AptaTaq DNA polymerase we used was already quite high at 15 Units per reaction, we kept the DNA polymerase concentration the same; increasing to 50 U did not further speed up the reaction (data not shown). In this thicker chamber and higher concentration primer reaction, we were able to observe threshold times of roughly 8 minutes, and the PCR reaction reached completion/saturation at 11 minutes (Fig. S32).

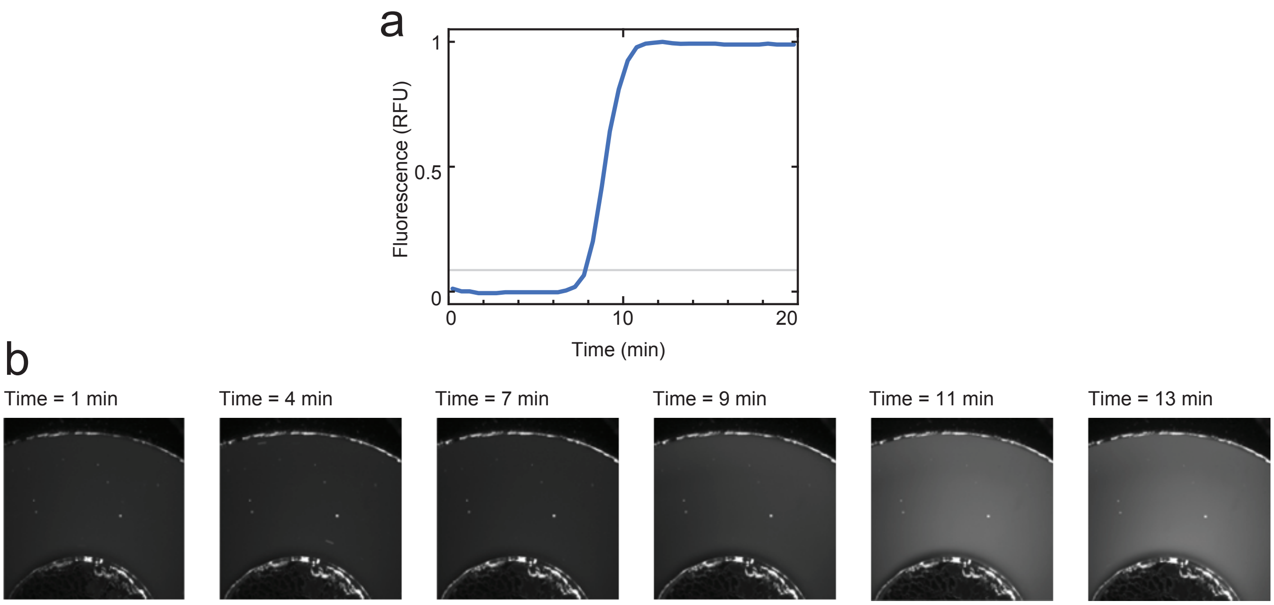


FIG. S32: **(a)** Ultrafast Donut PCR using a 700 $\mu$m-thick chamber and 2 $\mu$M primer concentration. The primers were designed to amplify the rs646776 SNP locus, and 10 ng of NA18537 human gDNA was used as input. Here, we added 0.82 $\mu$M of the SYTO-82 intercalating dye to visualize amplicon accumulation. As in other assays, the heaters of the amplification device were set to 60 °C and 95 °C. **(b)** Fluorescence images of the Donut PCR reaction at different timepoints. Note that no probes were printed in the chip for this reaction; instead, the bulk solution fluorescence was measured every 30 seconds for 20 minutes.

#### Feasibility of scaling to massively multiplexed panels

We believe that the Donut PCR platform has the capability to scale orders of magnitude past the 15- to 40-plex demonstrations in this manuscript. Over the past 2 decades, microarray fabrication technologies have advanced and now allows routine construction of custom microarrays with about 1 million probes. Although our Donut PCR probe array process will necessarily differ because we require three-stranded, fluorophore-functionalized DNA probes rather than single-stranded unmodified probes, the technologies for standard microarray can likely be adapted. In Fig. S33, we show a 33x33 printed probe array on a Donut PCR chip, unquenched to facilitate spot visualization. Out of the 1089 spots, only 1 pair of spots was unintentionally merged. Thus, the bacterial identification panel could easily be scaled to cover 1000 most frequent bacteria by printing additional 16S probes.

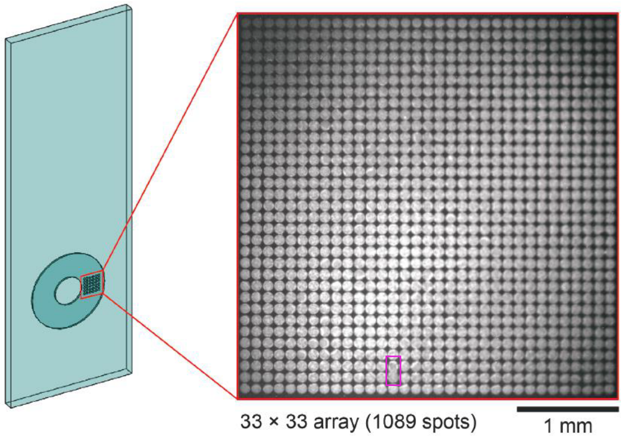


FIG. S33: A 33x33 array of multi-stranded probes printed using the same protocol described in the Materials and Methods, except using PDC-50 micro-nozzles. Only 1 pair of spots resulted in spot merging (purple), indicating that alignment and printing is mostly reproducible at these scales. Here, all spots contained probes of the same sequence; we expect similar results for printing an array of 1000 different probes.

[1] Ioannou, D., Huda, W., Laine, A.F. Circle recognition through a 2D Hough Transform and radius histogramming. *Image and Vision Computing*, 17, 15-26 (1999).
[2] D. Y. Zhang, S. X. Chen, P. Yin, Optimizing the specificity of nucleic acid hybridization. Nature chemistry, 4(3), 208-214 (2012).
